# Functional Interpretation of Genetic Variants Using Deep Learning Predicts Impact on Epigenome

**DOI:** 10.1101/389056

**Authors:** Gabriel E. Hoffman, Eric E. Schadt, Panos Roussos

## Abstract

Identifying causal variants underling disease risk and adoption of personalized medicine are currently limited by the challenge of interpreting the functional consequences of genetic variants. Predicting the functional effects of disease-associated protein-coding variants is increasingly routine. Yet the vast majority of risk variants are non-coding, and predicting the functional consequence and prioritizing variants for functional validation remains a major challenge. Here we develop a deep learning model to accurately predict locus-specific signals from four epigenetic assays using only DNA sequence as input. Given the predicted epigenetic signal from DNA sequence for the reference and alternative alleles at a given locus, we generate a score of the predicted epigenetic consequences for 438 million variants. These impact scores are assay-specific, are predictive of allele-specific transcription factor binding and are enriched for variants associated with gene expression and disease risk. Nucleotide-level functional consequence scores for non-coding variants can refine the mechanism of known causal variants, identify novel risk variants and prioritize downstream experiments.

## INTRODUCTION

Genome-wide association studies (GWAS) have identified thousands of loci associated with risk to human diseases (Visscher et al. 2017). Yet progress in understanding the molecular etiology of disease and the development of novel therapies has been limited by the fact that these studies are often not able to identify a specific causal variant and mechanistically relevant gene due to linkage disequilibrium (LD) (Visscher et al. 2017; Spain and Barrett 2015; Hindorff et al. 2009; Pritchard and Przeworski 2001). Integrating independent biological knowledge has the potential to increase the resolution of the associated region and improve the interpretation of GWAS results (Claussnitzer et al. 2015; Kichaev et al. 2014; Pickrell 2014). Most notably, risk variants are enriched in non-coding regulatory regions (Finucane et al. 2015; Maurano et al. 2012; Farh et al. 2015). While interpreting the functional consequences of protein coding variants has been remarkably successful and improved the understanding of the biology of human disease (Lek et al. 2016; Chong et al. 2015), the rules governing the functional effects of variants in non-coding regulatory DNA have been more challenging to decipher. Novel approaches are needed to interpret non-coding variants from ongoing whole genome sequencing projects, for example, of somatic variants in cancer (Zhang et al. 2018) and *de novo* variants in autism (Werling et al. 2018).

Recent work has sought to better understand the regulatory genome by characterizing the epigenetic differences in transcription factor (TF) binding, chromatin accessibility and histone modifications between tissues and cell types (Roadmap Epigenomics Consortium et al. 2015; ENCODE Project Consortium 2012; Andersson et al. 2014). Yet these epigenetic tracks can cover a substantial portion of the genome, even though polymorphisms at only a fraction of sites are presumed to have a functional consequence. Moreover, these efforts have generally not integrated genetic variation. Other efforts have focused on the effects of genetic variation on gene expression (Aguet et al. 2017; Lappalainen et al. 2013; Fromer et al. 2016) as well as multiple epigenetic assays (Grubert et al. 2015; Waszak et al. 2015; Chen et al. 2016a). Yet these xQTL studies are subject to the same challenges with linkage disequilibrium as GWAS so they generally cannot pinpoint the causal variant, or predict the functional consequence of a rare variant not observed in the dataset.

Recent progress in developing computational models able to predict TF binding, chromatin accessibility, and histone modifications from only the genome sequence in the surrounding region offers a novel paradigm to interpret the functional consequences of non-coding variants (Zhou and Troyanskaya 2015; Kelley et al. 2018; Lee et al. 2015; Kelley et al. 2016; Zhou et al. 2018). These models leverage advances in deep learning (LeCun et al. 2015) to use DNA sequence context in the predictive model of functional consequences. Yet with one exception (Kelley et al. 2018), these computational approaches consider only the discrete absence versus presence of an epigenetic signal (Zhou and Troyanskaya 2015; Lee et al. 2015; Kelley et al. 2016; Zhou et al. 2018). Moreover, these methods rely on the sequence of the reference genome so they do not model the contribution of genetic variation driving the epigenetic signal. Based on the extensive contribution of genetic variation to molecular phenotypes (Grubert et al. 2015; Waszak et al. 2015; Chen et al. 2016a; Aguet et al. 2017; Fromer et al. 2016; Lappalainen et al. 2013), and the increasing availability of epigenetics datasets from multiple individuals paired with genetic data (Girdhar et al. 2018; Waszak et al. 2015; Chen et al. 2016a; Grubert et al. 2015), integrating genetics into model training has the potential to improve prediction accuracy and increase power of variant impact predictions. Finally, although these methods are trained jointly across many datasets, they only consider a single experiment from a given cell type and assay.

Here we introduce a deep learning framework for <u>f</u>unctional <u>i</u>nterpretation of <u>g</u>enetic <u>v</u>ariants (DeepFIGV). We develop predictive models of quantitative epigenetic variation in chromatin accessibility from DNase-seq and histone modifications (H3K27ac, H3K4me3, and H3K4me1) from 75 lymphoblastoid cell lines (LCL) (Grubert et al. 2015). By modeling quantitative variation in the epigenetic signal, integrating whole genome sequencing to create a personalized genome sequence for each individual, and training the models on many experiments from the same cell type and assay, we identify variants with functional effects on the epigenome.

## RESULTS

### Deep learning maps from genome sequence to epigenetic signal

DeepFIGV combines the quantitative signal from epigenetic experiments across multiple individuals with whole genome sequencing into a single machine learning task (Figure 1). While standard xQTL analyses rely on the correlation between the epigenetic signal and a given genetic variant (Figure 1A-B), deep learning using a convolutional neural network explicitly models the DNA sequence context to train a predictive model (Figure 1C-D). Evaluating the predicted effect of each variant produces a large database of nucleotide-level scores (Figure 1E-F) that can be integrated with other analyses to refine the mechanism of known causal variants, identify novel risk variants and prioritize downstream experiments (Figure 1G).

**Figure 1:**
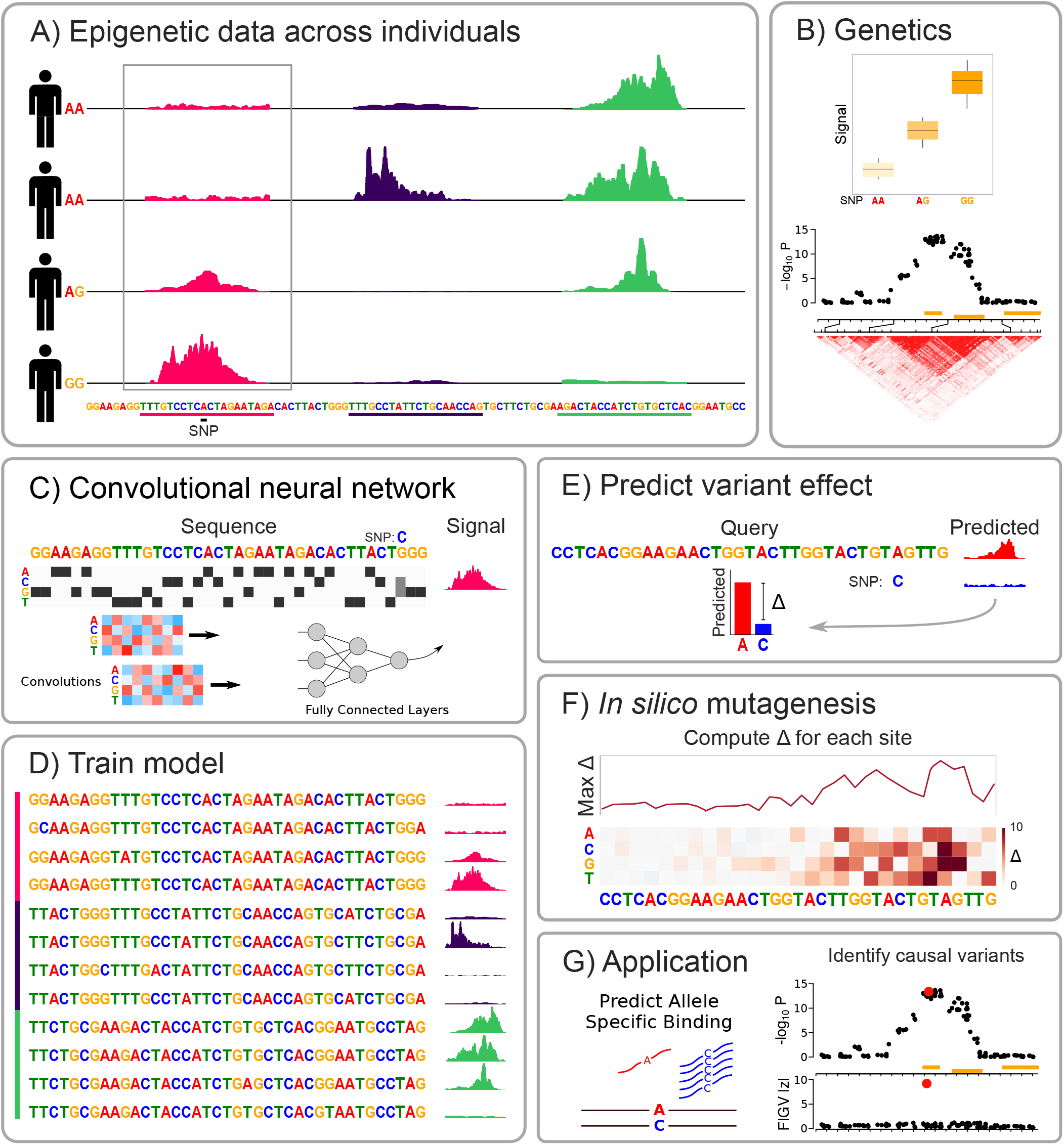
Computational workflow for Deep <u>F</u>unctional <u>I</u>nterpretation of <u>G</u>enetic <u>V</u>ariants (DeepFIGV). **A)** Quantitative signal from epigenetic assay (i.e. ChIP-seq, DNase-seq) across multiple individuals and genomic regions. **B)** Standard genetic analysis stratifies quantitative signal by the allelic state at a given SNP, yet linkage disequilibrium complicates the interpretation of the causal variant. **C)** DeepFIGV encodes a DNA sequences as an ‘image’ matrix of mostly zeros with a 1 (i.e. a dark box) indicating the presence of a particular nucleotide at that position. Heterozygous SNPs are encoded as a 0.5 for each allele. Convolutions are local matrix operations with parameter values learned from the data. A neural network uses the convolutions to predict the epigenetic signal from the DNA sequence. **D)** Training the computational model links DNA sequences from many individuals to the epigenetic signal in each region. **E)** The epigenetic signal is estimated for a query sequence with the reference and the alternate allele. The difference between the estimated signal values (i.e. delta) indicates the predicted effect of the variant. **F)** *In silico* mutagenesis evaluates the delta value for every possible single nucleotide substitution. **G)** DeepFIGV delta values are used to predict allele specific binding of transcription factors and identify candidate causal variants.

Datasets from each of four epigenetic assays (DNase-seq and H3K27ac, H3K4me3, and H3K4me1 histone marks) were analyzed separately (Figure 2). The parameters of the convolutional neural network were optimized to minimize the least squares prediction error. Extensive steps were taken to avoid overfitting, and all prediction results are reported on a set of individuals and chromosomes that were excluded from the training set (see Methods, Supplementary Figure 1). Increasing the number of individuals in the training set and including genetic variation in the genome sequence of each individual decreased prediction error on withheld test data (Supplementary Figure 2). Although the model uses only DNA sequence in the predictions, the predicted DNase signal shows strong concordance in the test set with the observed signal (Spearman rho = 0.485, Pearson R = 0.707) (Figure 2A). Focusing on the more robust (i.e. rank based) Spearman correlation metric shows that these predictive models give substantial accuracy for all four assays for the quantitative epigenetic signal (Figure 2B). Examining the predicted signal for all four assays from a representative example in the test set along a segment of chromosome 22 shows notable concordance with the observed signal (Figure 2C).

**Figure 2:**
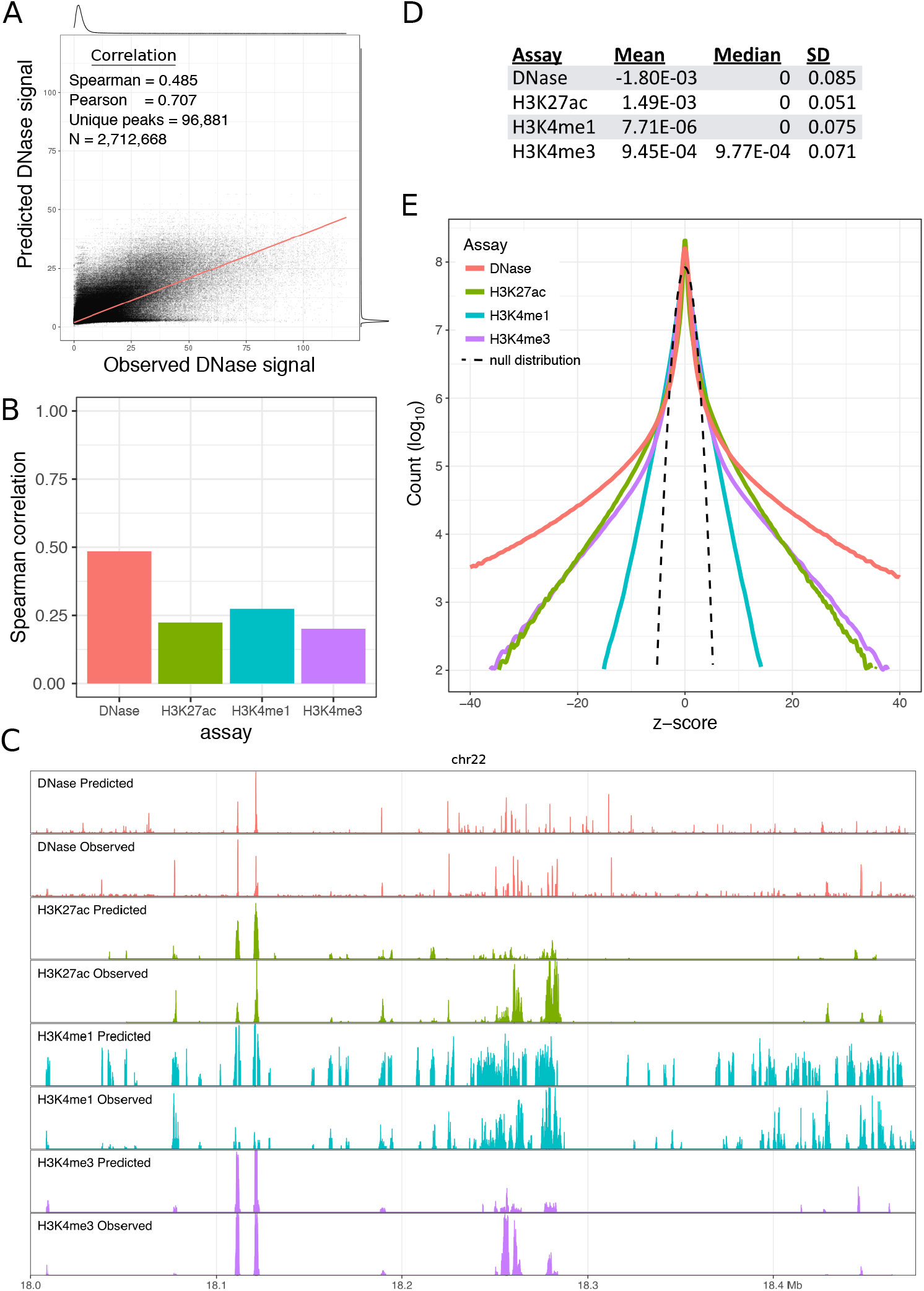
Evaluating DeepFIGV model and interpreting variant scores. **A)** Predicted DNase signal compared to observed DNase signal evaluated on the test set. **B)** Spearman correlation between predicted and observed epigenetic signal on the test set for 4 assays. **C)** Predicted and observed DNase signal for a 300kb segment on chr22 in the test set. **D)** Mean, median and standard deviation of the delta scores for 208 million biallelic SNPs for each assay. **E)** Density plot of z-scores for 4 assays. Dashed line indicates the null distribution of the z-scores, which is the standard normal distribution.

Functional impact scores from the predicted difference in epigenetic signal for the reference versus the alternate allele were evaluated for 208 million biallelic SNPs from the gnomAD database of 15 thousand whole genome sequences (Lek et al. 2016). The delta value for each variant is defined as Δ = S_ALT_ – S_REF_ with terms representing the predicted epigenetic signal from the alternative and reference alleles, respectively. Thus, a positive delta value indicates that the alternative allele increases the epigenetic signal compared to the reference allele. As expected, the mean and median delta values for all assays were very close to zero (Figure 2D). Transforming these delta values to a standard scale by dividing by the standard deviation for each assay shows an excess of variants with scores near zero compared to the standard normal distribution (Figure 2E). This is consistent with the vast majority of variants having no functional effect on the epigenome. Yet there is an excess of variants with large effects on all four epigenetic assays, with DNase showing the highest excess followed by H3K27ac, H3K4me3 and finally H3K4me1.

### Genomic correlates of predicted variant effects

Although no prior biological information is included in training the model, DeepFIGV recovers multiple aspects of known regulatory biology (Figure 3). The predictive model learned by the convolutional neural network is composed of a set of local sequence features called filters. Although learned *de novo*, the predictive sequences features extracted by these filters are often similar to known transcription factor bindings site (TFBS) motifs. Some filters have a direct correspondence to a known motif, but other filters model only a portion of a motif so that multiple filters combine to capture the signal encoded by the sequence (Figure 3A). Variants in TFBS motifs are enriched for the alternative allele decreasing the DNase signal (Figure 3B). TFBS nucleotides with high information content (i.e. high weight) in the position weight matrix have an even stronger enrichment for decreasing the DNase signal, consistent with variants being more likely to weaken rather than strengthen the affinity of a TFBS motif (Supplementary Figure 3). The TFBS enrichments are consistent with the biology of these assays: variants predicted to affect the open chromatin assay DNase are most enriched for TFBS motifs, followed by the H3K4me3 promoter mark and the H3K27ac active promoter and enhancer mark (Figure 3C). H3K4me1 is an enhancer mark that tags active or repressed sequences and is not enriched for strong variants in TFBS.

**Figure 3:**
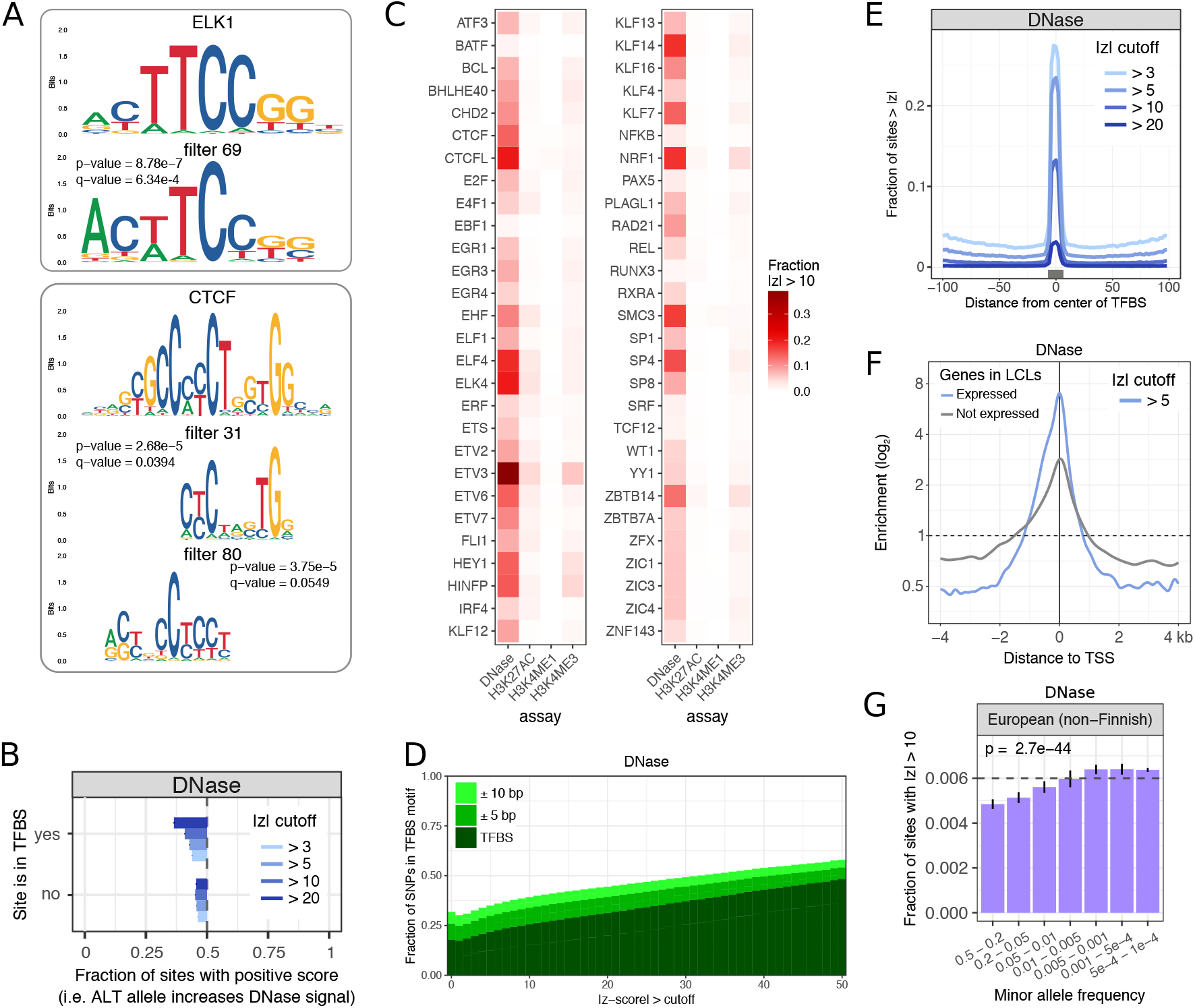
Genomic enrichments of predicted functional variants. **A)** Canonical transcription factor binding motif along with the motif representation of convolutional filters learn from the DNase dataset. P-values indicate the probability of concordance this high between a canonical motif and convolutional filter occurring by chance given the motif database. Q-values correct for multiple testing since 300 convolutional filters were queried. **B)** Ratio indicating the fraction of sites where the alternative allele increases the DNase signal (i.e has a positive DeepFIGV z-score) for DNase for 4 cutoffs. Ratios are shown for variants within or outside a TFBS motif. A value of 0.5 indicates an equal number of variants with positive and negative z-scores. A value < 0.5 indicates an depletion of variants where the alternative allele increases the DNase signal, corresponding to an excess of variants where the alterative allele decreases DNase signal. **C)** Fraction of variants in transcription factor binding sites that exceed a DeepFIGV absolute z-score of 10 for each of 4 assays. **D)** Fraction of sites that are in a transcription factor binding site motif, or in the flanking 5 or 10 bp, for a range of DeepFIGV absolute z-score cutoffs for DNase. **E)** Enrichment of variants near a TFBS motif exceeding 4 z-score cutoffs for DNase. Black box indicates median size of TFBS motif. **F)** Enrichment of sites with absolute z-scores greater than 5 near the transcription start site of genes stratified by whether the genes are expressed in LCLs. Sites with absolute z-scores less than the genome-wide mean are used as the baseline for the enrichment. **G)** Fraction of sites with absolute z-scores for DNase greater than 10 within 7 minor allele frequency bins based on non-Finnish Europeans from gnomAD. Dashed line indicates genome-wide fraction of sites. P-value is based a logistic regression where the response is a binary variable indicating if the absolute z-scores is greater than 10 and the log minor allele frequency is the predictor. Error bars show 95% confidence intervals.

The role of TFBS in regulating gene expression and epigenetics is well established, and consequences of variants in TFBS motifs are more interpretable than variants in other genome annotations (ENCODE Project Consortium 2012; Kheradpour and Kellis 2014; Lambert et al. 2018). Yet despite notable enrichment in TFBS motifs, variants in these motifs account for a minority of variants with strong DeepFIGV scores. Only 15.2% of sites with an absolute z-score between 9 and 10, and 19.9% with an absolute z-score between 19 and 20 for DNase fall in a TFBS (Figure 3D, Supplementary Figure 4). Including flanking nucleotides within 5 or 10 bp increases these percentages, but for most z-score cutoffs, variants in or proximal to known TFBS motifs are a minority. While variants in TFBS motifs are enriched for variants predicted to affect DNase signal, the enrichment is not observed when expanding beyond these proximal nucleotides (Figure 3E for DNase, Supplementary Figure 3C for ChIPseq). Therefore, the majority of variants with strong predicted effects on all four assays do not fall in nor are they proximal to these known TFBS, indicating that DeepFIGV models a more complicated relationship between genetic variants in epigenetic signal than is encoded by TFBS motifs alone.

Variant effects show a degree of cell type specificity as variants with strong predicted effects on DNase, H3K27ac and H3K4me3 are more enriched around the transcription start site (TSS) of genes expressed in LCLs, compared to genes not expressed in LCLs (Figure 3F, Supplementary Figure 5). Variants with strong predicted effects are also more enriched around the TSS of LCL-specific genes, compared to tissue-specific genes from each of 52 additional GTEx tissues (Supplementary Figure 6). Moreover, variants with strong predicted effects on DNase, H3K27ac and H3K4me3 are enriched in CpG islands and ChromHMM tracks from LCLs (Supplementary Figures 7 and 8). Finally, variants with large predicted effects are depleted among common variants (minor allele frequency > 1%) and are enriched in rare variants across multiple human populations, consistent with negative selection against variants that disrupt the epigenome (Figure 3G, Supplementary Figure 9).

### Concordance with xQTLs

Lead cis-QTL variants (i.e. the local variant with the smallest p-value) for multiple assays are enriched for having strong predicted effect on the epigenome (Figure 4, Supplementary Figure 10). Variants that are lead cis-QTLs for DNase from the current dataset (Grubert et al. 2015) are particularly enriched for having a strong predicted effect on DNase and H3K4me1 (Figure 4A). Similarly, variants that are lead cis-QTLs for gene expression in an independent dataset of LCLs from European individuals (Lappalainen et al. 2013) are most enriched for variants with a strong predicted effect on DNase (Figure 4B). Rare variants associated with expression outliers in multiple tissues types (Li et al. 2017a) are enriched for variants with strong predicted effects on DNase (Figure 4C), but not ChIP-seq. Moreover, somatic variants in cancer that drive changes in expression of nearby genes are enriched for variants with strong predicted effects on DNase, H3K4me3 and H3K27ac (Zhang et al. 2018) (Figure 4D). Furthermore, candidate causal variants for expression QTLs identified by statistical fine mapping (Brown et al. 2017) are enriched for variants with strong predicted effects on DNase (Figure 4E). For example, rs11547207 is identified as an eQTL in LCLs from European individuals, but this SNP is in linkage disequilibrium with many nearby variants (Figure 4F). Statistical fine mapping indicates that this SNP has a high probably of being the causal variant in this region driving gene expression variation. Although analysis of the DNase signal from LCLs does not identify this SNP as a QTL, DeepFIGV directly models the sequence context of this variant and predicted a strong effect of the epigenetic signal in this region. *In silico* saturation mutagenesis in this region gives predictions at nucleotide-resolution and indicates that variants within ~5 bp are also predicted to decrease the DNase signal despite not falling in a known TFBS (Kheradpour and Kellis 2014).

**Figure 4:**
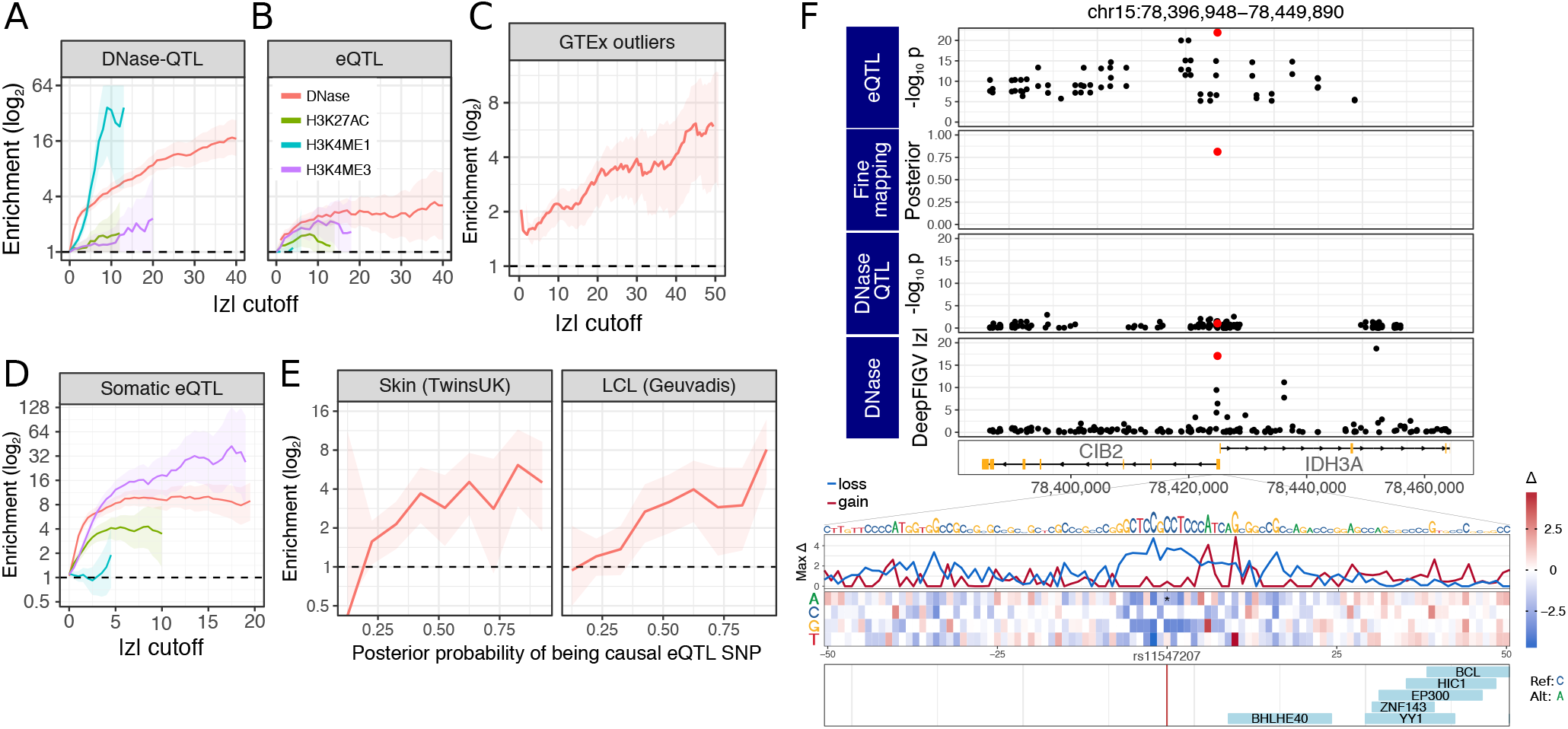
DeepFIGV scores predict results of xQTL analysis. **A,B)** Lead variants from xQTL analysis from lymphoblastoid cell lines are enriched for SNPs with DeepFIGV absolute z-score exceeding a range of cutoffs. Enrichments are evaluated using **(A)** DeepFIGV scores for 4 assays for DNase-QTLs and **(B)** expression QTLs. Shaded regions indicated 95% confidence intervals. **C)** Rare variants associated with gene expression outliers are enriched for DeepFIGV absolute z-score for DNase compared to rare variants not associated with outliers. Shaded regions indicated 95% confidence intervals. **D)** Enrichment of somatic variants in cancer that drive gene expression changes (Zhang et al. 2018) for strong DeepFIGV scores. **E)** Candidate causal variants for expression QTLs with higher posterior probability are enriched for exceeding a DeepFIGV absolute z-score of 10 for DNase. Enrichments are shown for skin and LCL samples from TwinsUK, and LCL samples from GEAUVIDIS (Lappalainen et al. 2013). Shaded regions indicated 95% confidence intervals. **F)** DeepFIGV independently identifies candidate causal variant rs11547207 (shown in red) for QTL affecting expression of both CIB2 and IDH3A. eQTL analysis of GEUAVDIS identifies many correlated variants associated with these genes, but statistical fine-mapping identifies a signal candidate variant (Brown et al. 2017). Although this variant is not an DNase QTL in LCLs Yoruban individuals (Grubert et al. 2015), DeepFIGV analysis on the same data identifies the same candidate causal variant identified by statistical fine mapping. *In silico* mutagenesis of 50bp around rs11547207 indicates that variants at nearby positions are predicted to decrease the DNase signal. Size of letters in DNA sequence is proportional to the maximum absolute delta at that position. Bottom panel shows TFBS motifs.

### DeepFIGV variant scores predict allele specific binding

The predicted functional effect of genetic variants on each of the four epigenetic assays analyzed in DeepFIGV can identify allele-specific binding (ASB) of TFs in independent ChIP-seq experiments in LCLs (Chen et al. 2016b; Shi et al. 2016) (Figure 5). Heterozygous variants can be divided into 3 categories based on ASB: no allele specific effect, ASB favoring the reference allele, and ASB favoring the alternative allele (Figure 5A). We evaluated the ability of DeepFIGV to distinguish between these categories even though no allele-specific information is included in model training. The predicted effect on the DNase signal can classify variants showing ASB versus no ASB for CCCTC-binding factor (CTCF) with an area under the precision recall (AUPR) curve of 0.202 compared to 0.0493 for a random classifier (Figure 5B,C). Given a variant with an allele-specific effect, the predicted effect on DNase signal is able to classify the direction of the effect (i.e. favoring reference versus alternative) for CTCF with an AUPR of 0.704 compared to a random classifier of 0.36 (Figure 5D,E). Since the number of sites in each category varied substantially across TFs, we consider the increase in AUPR from the DeepFIGV score compared to a TF-specific baseline. DeepFIGV scores show an increase in AUPR compared to a TF-specific random classifier for distinguishing the ASB status of variants for independent assays of multiple transcription factors in LCLs (Figure 5F) and HeLa S3 cells (Supplementary Figure 11).

**Figure 5:**
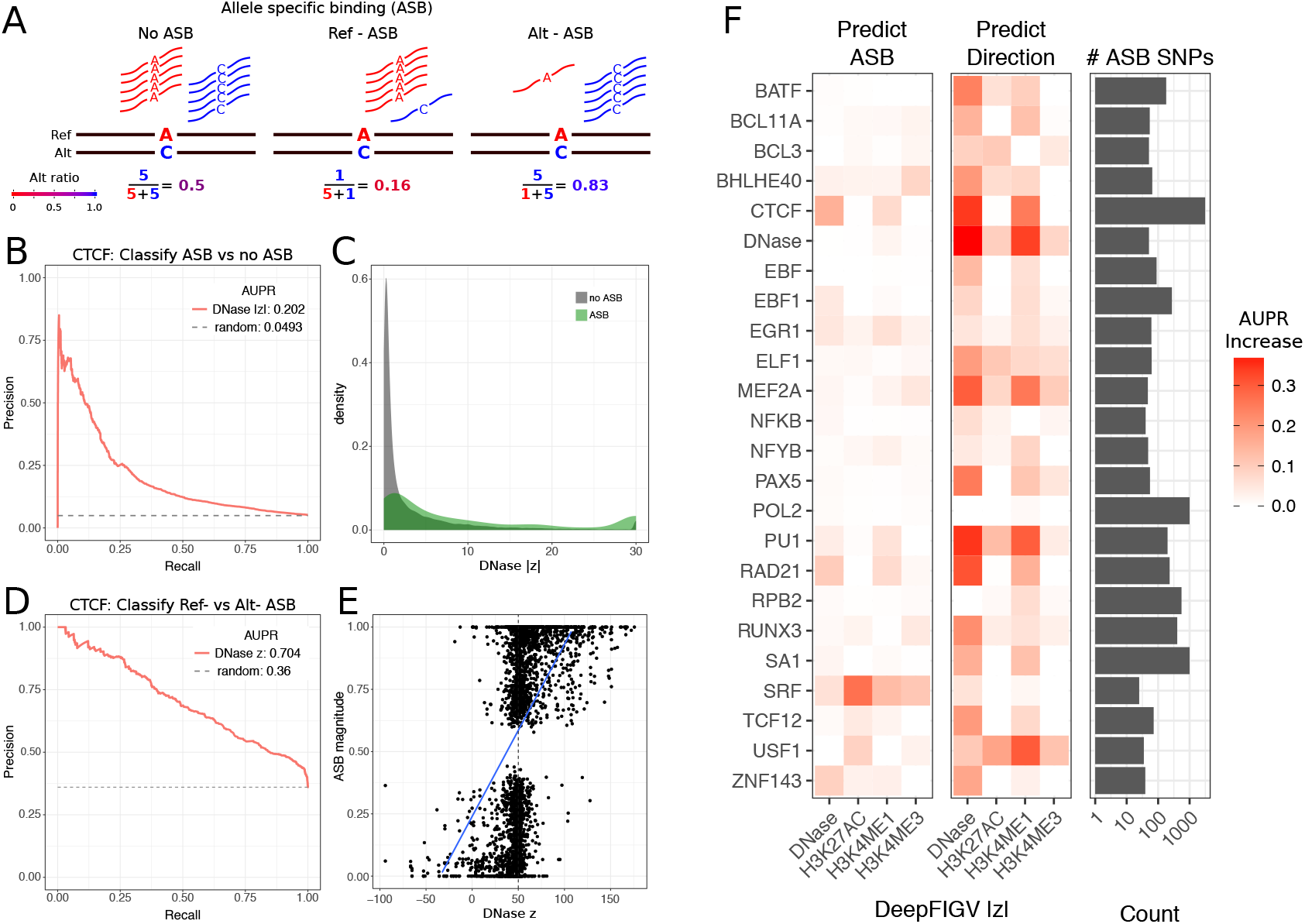
DeepFIGV scores predict allele specific transcription factor binding in LCLs. **A)** Diagram illustrating 3 categories of allele specific binding (ASB): 1) no ASB, 2) ASB favoring the reference allele, and 3) ASB favoring the alternative allele. **B)** Precision-recall curve indicating performance of absolute DeepFIGV z-score for DNase in predicting ASB of CTCF. AUPR indicates the area under the precision-recall curve. Dashed line indicates the performance of a random predictor. **C)** Density plot showing absolute DeepFIGV z-score for variants in **(B)** in the ABS or no ASB classes. **D)** Precision-recall curve indicating performance of DeepFIGV z-score for DNase in predicting the directionality of ASB (reference versus alternative) for CTCF. AUPR indicates the area under the precision-recall curve. Dashed line indicates the performance of a random predictor. **E)** Plot of ASB magnitude versus DeepFIGV DNase z-score from **(D). F)** Increase in AUPR of predicting ASB status for DeepFIGV scores for 4 epigenetic assays compared to a TF-specific random predictor. Increase in AUPR is shown for predicting ASB versus no ASB (left) and predicting the directionality of ASB (reference versus alternative) (center). Right panel shows the number of ASB SNPs considered in each analysis.

### Enrichment for disease risk variants and interpreting causal variants

Integrating DeepFIGV scores with large-scale genome-wide association studies shows that risk variants for common disease are enriched for variants predicted to impact the epigenome (Figure 6). We applied stratified LD-score regression (Finucane et al. 2015) to evaluate the contribution of variants with different genomic annotations to disease risk. Analysis of 19 traits identified a contribution of variants passing multiple DeepFIGV z-score cutoffs to trait heritability, even after accounting for a baseline set of 32 genomic annotations (see Methods) (Figure 6A, Supplementary Figure 12,13). Immune traits show the strongest contribution of DeepFIGV variants to trait heritability since the model was trained in LCLs (a B-cell lineage), yet there are also cell type autonomous effects and a contribution to non-immune traits. Further investigation of the impact of immune traits shows that candidate causal variants identified by statistical fine mapping (Farh et al. 2015) are enriched for variants with strong DeepFIGV effects (Figure 6B, Supplementary Figure 14).

**Figure 6:**
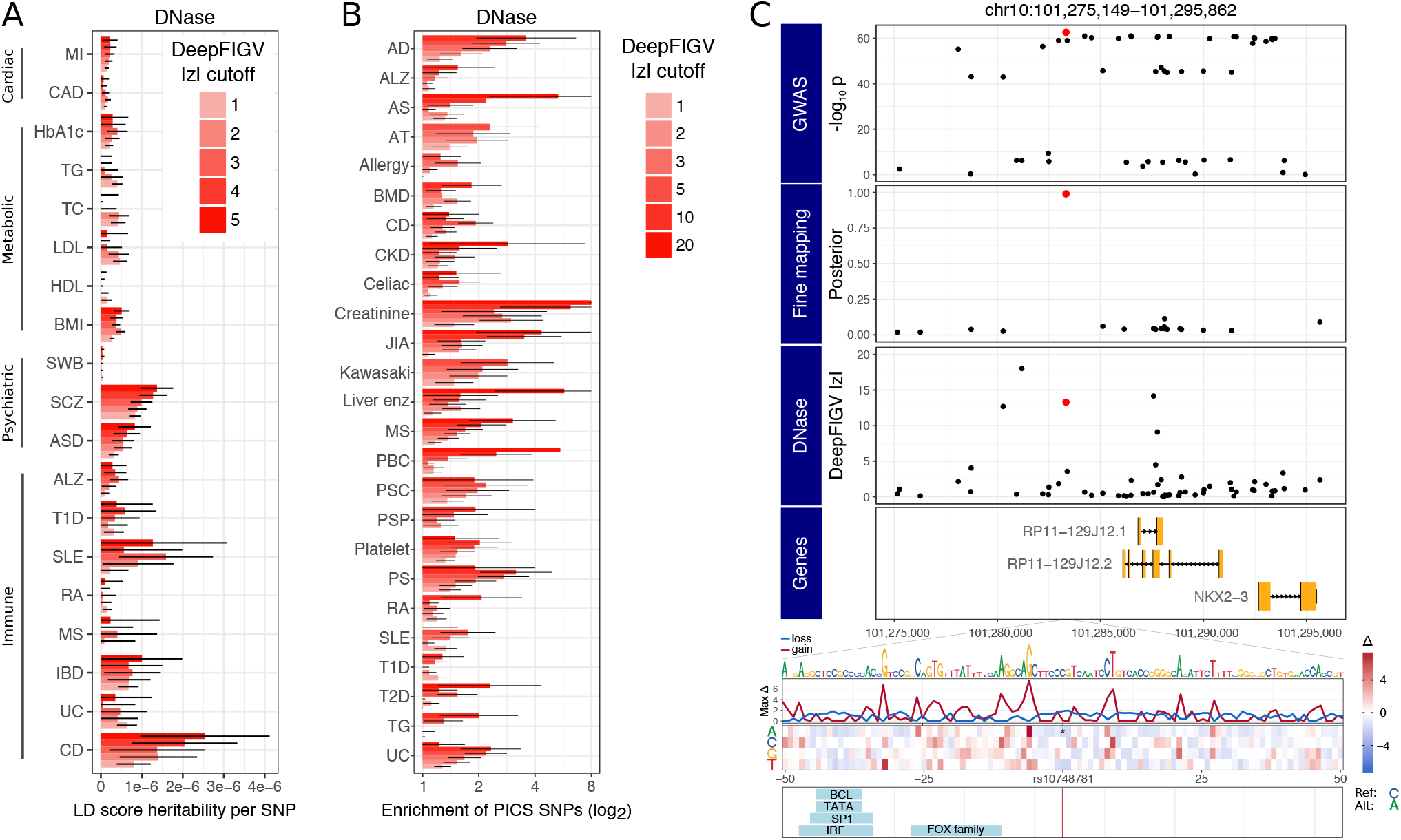
Disease risk variants are enriched for large DeepFIGV scores. **A)** Linkage-disequilibrium score regression (LDSC) (Finucane et al. 2015) partitioned heritability estimates for diseases in 4 categories. Heritability per SNP is computed for variants that exceed 5 cutoffs for DeepFIGV absolute z-score for DNase. Error bars indicate 2 standard deviations. **B)** Enrichment of candidate causal variants for autoimmune disease (Farh et al. 2015) are variants exceeding 6 cutoffs DeepFIGV absolute z-score for DNase. Error bars indicate 2 standard deviations. **C)** DeepFIGV elucidates molecular function of candidate causal variant for inflammatory bowel disease (Huang et al. 2017a). GWAS identifies many correlated variants associated with disease risk, but statistical fine mapping identifies a single SNP (shown in red) as the candidate causal variant. This variant, rs10748781, disrupts a CpG site and is predicted to decrease the DNase signal in this region. *In silico* mutagenesis of 50bp around this SNP indicates that variants at nearby positions are predicted to decrease the DNase signal. Size of letters in DNA sequence is proportional to the maximum absolute delta at that position. Bottom panel shows TFBS motifs. Disease abbreviations: AD (Atopic dermatitis), ALZ (Alzheimer’s), AS (Ankylosing spondylitis), ASD (Autism spectrum disorder), AT (Autoimmune thyroiditis), BMD (Bone mineral density), BMI (Body mass index), CAD (Coronary artery disease), CD (Crohn’s disease), CKD (Chronic kidney disease) HbA1c (HbA1c protein level in blood), HDL (High-density lipoprotein), IBD (Inflammatory bowel disease), JIA (Juvenile idiopathic arthritis), LDL (Low-density lipoprotein), Liver enz (gamma glutamyl transferase), MI (myocardial infarction), MS (Multiple sclerosis), PBC (Primary biliary cirrhosis), PSC (Primary sclerosing cholangitis), PSP (Progressive supranuclear palsy), PS (Psoriasis), RA (Rheumatoid arthritis), SLE (Systemic lupus erythematosus), SWB (Subjective well-being), T1D (Type 1 diabetes), T2D (Type 2 diabetes), TC (total cholesterol), TG (Triglycerides), UC (Ulcerative colitis).

DeepFIGV scores can elucidate the molecular mechanism of a causal variant and prioritize downstream experiments. Integrating DeepFIGV scores with candidate causal variants for inflammatory bowel disease (Huang et al. 2017a) shows that for rs10748781, which has 99% posterior probability of being the causal variant in this region, the alternative allele is predicted to decrease the DNase signal in this region in LCLs (Figure 6C). This result gives a specific cell type and biological assay to design a validation experiment. Moreover, this variant disrupts a CpG site, is a known DNA methylation QTL, and the methylation at nearby sites is predicted to mediate the effect of the variant on disease risk (Hannon et al. 2017) (Supplementary Figure 15).

## DISCUSSION

Translating findings of genetic studies to a molecular understanding of disease etiology and then to novel therapies has been hindered by the challenge of interpreting the functional consequence of genetic variants. There is a widely recognized need for accurate computational predictions of the functional impact of non-coding regulatory variants (Albert and Kruglyak 2015). Genomic annotations of the non-coding regions have generally taken one of four approaches. Evolutionary conservation or selection can identify functional regions of the genome, but consecutive nucleotides often have very similar scores and this approach does not give cell type- and assays-specific functional consequences (Siepel et al. 2005; Gulko et al. 2015; Huang et al. 2017b). Epigenetic maps across multiple cell types, tissues and assays provide a functional interpretation, but peaks from these assays cover millions of nucleotides (ENCODE Project Consortium 2012; Roadmap Epigenomics Consortium et al. 2015). xQTL studies correlate genetic variants with gene expression or epigenetic signals, but interpretation of this correlation analysis is limited by linkage disequilibrium and is only applicable to variants observed in the dataset (Aguet et al. 2017; Lappalainen et al. 2013; Fromer et al. 2016). Most recently, deep convolutional neural networks have been used to develop predictive models linking the genome sequence to splicing (Xiong et al. 2015), protein binding (Alipanahi et al. 2015), and epigenetic signals (Zhou and Troyanskaya 2015; Lee et al. 2015; Kelley et al. 2016; Zhou et al. 2018). Although these deep learning methods have been promising, their biological application has so far been limited.

Here we present a deep learning framework that learns a predictive model linking DNA sequence to quantitative variation in epigenetic signal and evaluates the predicted functional impact of genetic variants on multiple assays. This framework models quantitative variation in the epigenome, integrates whole genome sequencing to create a personalized genome sequence for each individual, and trains on many experiments from the same cell type and assay. Because this framework fits a predictive model based on sequence context, it is less susceptible to issues of linkage disequilibrium and can predict the functional impact of variants even if they are not observed in the training dataset.

Application to epigenetic assays of open chromatin (DNase-seq) and histone modifications (H3K27ac, H3K4me3, and H3K4me1) from 75 lymphoblastoid cell lines (LCL) (Grubert et al. 2015) produces functional consequence scores that are concordant with other genomic annotations while capturing sequence context information beyond known TFBS motifs. We note that potential mechanisms of variants outside these motifs include affecting local DNA shape, DNA methylation or nucleosome positioning (Slattery et al. 2014; Deplancke et al. 2016), but interpretation remains an open challenge (Shrikumar et al. 2017). We demonstrate that these functional consequence scores inform molecular mechanism, are concordant with xQTL analysis, are predictive of allele-specific binding, and inform interpretation of risk variants for common disease. Moreover, these scores can prioritize variants for downstream experiments and indicate the appropriate cell type and functional assay. DeepFIGV scores are complementary to other non-coding variant scores, and compared to DeepSea (Zhou and Troyanskaya 2015) identifies more variants with extreme z-scores particularly for sites with minor allele frequency less than 5% (Supplementary Methods, Supplementary Figures 16,17).

The differing performance of the prediction and biological enrichments across the four epigenetic assays is attributable to both biological and technical factors. These assays differ in the biological processes they measure. DNase measures open chromatin with high signal representing protein interacting with the DNA within a narrow region of ~150 bp. Thus DNase signal is largely determined by the proximal DNA sequence and especially TF binding. Histone modification ChIP-seq is more complex readout of chromatin state with H3K4me3 at active promoters, H3K27ac at active promoters and enhancers, and H3K4me1 at either active or repressed enhancers. Due to spatial chromatin spreading, the role of trans-factors, and the increased width of these marks (300 bp to 1kb), sequence-based prediction is known to be less accurate (Zhou and Troyanskaya 2015). Since genetic variants conferring disease risk or regulating gene expression can act through a number of mechanisms (Grubert et al. 2015; Maurano et al. 2012; Huang et al. 2017a; Aguet et al. 2017; Albert and Kruglyak 2015), the value of additional epigenetic assays depends on the accuracy of a predictive model as well as the regulatory mechanism of interest.

Scalable experimental approaches to measure the functional consequence of non-coding variants have recently been proposed (Starita et al. 2017; Tewhey et al. 2016; Ulirsch et al. 2016; Ernst et al. 2016; Arnold et al. 2013). These massively parallel report assays (MPRA) couple thousands to millions of nucleotide sequences to a molecular readout that can be quantified by short read sequencing. Despite their remarkable experimental throughput, these assays are limited to cell culture and they assay the function of the query sequence either in an episomal vector or through random insertion into the genome. Thus the degree to which results from MPRAs recapitulate function in the disease relevant cell type and natural genomic context remains unclear (Inoue et al. 2017; Muerdter et al. 2018; Ernst et al. 2016). In contrast, predictive models based on sequence context use natural genetic variation, are extensible to multiple biological assays, and evaluate sequences in their native chromosomal context. Moreover, they are applicable to cell culture, as well as cells from post mortem, biopsy or blood draws to more precisely target the relevant cell type. In fact, variants found to drive changes in gene expression in LCLs by a recent MPRA (Tewhey et al. 2016) are enriched for having strong predicted effects on DNase by DeepFIGV (Supplementary Figure 18).

The growth of large-scale resources pairing quantitative epigenetic assays with genetic data offers an opportunity to train rich predictive models on disease relevant cell types (PsychENCODE Consortium et al. 2015; Chen et al. 2016a; Girdhar et al. 2018). Finally, we have developed a public resource of the DeepFIGV predicted functional scores for 438 million variants available at <u>deepfigv.mssm.edu</u>.

## METHODS

### Epigenomic data from lymphoblastoid cell lines

The dataset comprises ChIP-seq experiments for 3 histone modifications (H3K27ac, H3K4me1 and H3K4me3) for 75 individuals and DNase I hypersensitivity experiments for 69 individuals (Grubert et al. 2015). All individuals are of Yoruban ancestry from the 1000 Genomes Project (The 1000 Genomes Project Consortium 2012). Processed data was downloaded from the ChromoVar3D website (<u>chromovar3d.stanford.edu</u>). Peak coordinates and signal intensities for each sample and each DNase peak were extracted from `DNase_removeBlacklist_Log10PvalueThreshold_5_DATA_MATRIX.gz`, and corresponding files were used for the 3 histone modifications. VCF of variants from whole genome sequencing was obtained from the same website.

### Deep learning with a convolutional neural network

The basset software (Kelley et al. 2016) was used to learn parameters in a predictive model mapping from genome sequence as input to epigenetic signal as output. The analysis was customized to take advantage of this particular dataset by 1) integrating genetic variation from whole genome sequencing, 2) modeling the quantitative variation in the epigenetic signal and 3) combining many experiments from the same cell type into a large single-task learning application. This customized analysis enables a focus on genetic variants with relatively small effects on the quantitative signal value, rather than the strong effect required to completely lose or gain a binding or histone modification event. Each of the 4 assays was analyzed separately using a single-task learning approach.

### Constructing DNA sequence as input to neural network

Personalized genome sequences were constructed using the GRCh37 reference genome with sites modified according to biallelic SNPs in the whole genome sequence using the `bcftools consensus` command. At homozygous alternate sites the reference allele is simply replaced by the alternative allele. Heterozygous sites are represented using the IUPAC nucleotide ambiguity codes (Comnish-Bowden 1985), so that for example an A/C heterozygote is indicated with the letter ‘M’. Only biallelic SNPs are considered, so there are 6 ambiguity codes, one for each pair of nucleotides.

Non-ambiguous sites are one-hot coded as a matrix of mostly 0’s with 4 rows corresponding to ‘A’, ‘T’, ‘C’, and ‘G’. Coding a 1 in the ‘T’ row indicates the presence of that nucleotide in the corresponding position in the genome sequence. Ambiguous sites are encoded with a value of 0.5 in the two corresponding rows. Thus the training data does not explicitly include any information about phasing of the SNPs or allele-specific signals.

For each peak interval called in the processed data, the genome sequence within a specified distance from the center of the peak was extracted and matched to the corresponding signal value. Peaks exceeding an assay-specific width cutoff were excluded from the analysis (Supplementary Table 1). An assay-specific window size between 300 and 2000 bp was used to extract regions from the personalized reference genomes (Supplementary Table 1). Larger window sizes have been shown to increase prediction performance (Zhou and Troyanskaya 2015), and we used the largest window size where encoding the DNA sequence from all peaks and all individuals to one-hot coded format could be computed on a machine with 256 Gb RAM.

### Model training and testing

Basset was trained with a 3 layer deep neural network and 300 convolutional filters each 19 bp wide (Supplementary Table 3). Changing the number of filters between 100 and 400 and changing the filter width between 10 and 20 bp did not produce a substantial change in prediction accuracy. A 30% dropout was applied to avoid overfitting. All training was performed on NVIDIA Tesla K20X GPU. Training on a single assay took between 14 and 45 GPU hours. Multiple restarts gave similar prediction accuracy.

The dataset was divided into training, validation and testing sets. In order to avoid overfitting, an early stopping approach was used where the parameter values in the model were learned from the training set, but the final values were selected to minimize the squared prediction error in the validation set. The prediction performance for each assay was reported based on the test set.

The training, validation and testing sets were specially constructed using a conservative approach in order to ensure independence of the three sets. Since the signal values at a given peak are relatively similar across individuals, including the same peak region, albeit from different individuals, in both the training and testing sets could overstate the prediction performance. Similarly, peaks from the same individual are generated under the same experimental conditions and are subject to technical batch effects. Thus, including peaks from the same individual in both the training and testing set could also overstate the prediction performance. In order to avoid this issue, the test set is composed of peaks on chr1-chr8 from 60% of individuals, the validation set is composed of peaks on chr9-chr15 from the next 20% of individuals, and the test set is composed of peaks on chr16-chr22 from last 20% of individuals. Thus, the three sets have no overlap in either peaks or individuals (Supplementary Table 1, Supplementary Figure 1) to ensure a conservative estimate of prediction performance.

We describe the analysis workflow with numbers from DNase data; numbers for other assays are shown in Supplementary Table 1. There were 681,990 total DNase peak intervals from 69 individuals. In order to focus on peaks of approximately equal size, peaks exceeding 250 bp were excluded. This left 463,094 peaks (67.9% of total) with a mean width of 150.7 bp. Multiplying the number of remaining peaks by the number of individuals gives a dataset of 31,953,486 examples. Since DNase and histone modification ChIP-seq are not strand specific-assays, the reverse complement sequence gives the same epigenetic signal as the original sequence.

Augmenting the dataset by including the reverse complement of each example doubles the number of sequence-signal pairs. Constructing the training set from peaks on chr1-chr8 from the first 60% of individuals gives 229,421 unique peak regions and 18,812,522 total examples.

### Genomic correlates

Minor allele frequency across populations were obtained from gnomAD r2.0.2 (Lek et al. 2016). Genomic locations of transcription factor binding motifs were obtained from ENCODE (Kheradpour and Kellis 2014). Genomic locations from ChIP-seq experiments for transcription factors in LCL GM12878 (ENCODE Project Consortium 2012) were downloaded from http://egg2.wustl.edu/roadmap/src/chromHMM/bin/COORDS/hg19/TFBS/gm12878/. List of genes expressed in LCLs were obtained from http://egg2.wustl.edu/roadmap/src/chromHMM/bin/COORDS/hg19/expr/gm12878/. ChromHMM tracks (Roadmap Epigenomics Consortium et al. 2015) for LCL GM12878 were downloaded from http://egg2.wustl.edu/roadmap/data/byFileType/chromhmmSegmentations/ChmmModels/core_K27ac/jointModel/final/E116_18_core_K27ac_dense.bed.gz. Genome annotation of sites were obtained from VEP v85 (McLaren et al. 2016) provided by gnomAD (Lek et al. 2016). CpG islands were obtained from Annotatr (Cavalcante and Sartor 2017).

### Comparison to canonical motifs

Filters learned by basset were compared to known motifs from CisBP (Weirauch et al. 2014) using tomtom (Gupta et al. 2007). Motifs were visualized using ggseqlogo (Wagih 2017).

### Evaluating variant effects

Coordinates and alleles of SNPs were obtained from multiple public resources (Supplementary Table 2) and combined into a non-redundant list comprising 413,223,060 sites and 437,960,283 variants (due to multi-allelic sites). The delta between the predicted signal from the reference and alternative alleles was evaluated for each of the four epigenetic assays. The median and standard deviation of the delta values for each assay were obtained for 208 million biallelic SNVs from whole genome sequencing (WGS) from gnomAD r2.0.2 and were used to compute z-scores for the entire set of variants for the corresponding assay. This approach used sites distributed across the genome that were identified independently of their predicted functional consequence and avoids double counting multiallelic sites. The standard deviation was computed using a robust method (i.e. winsorized) where delta values below the 1^st^ percentile or above the 99^th^ percentile were set to the value at the corresponding cutoff. This approach reduced the effect of variants with extreme scores. Changing the cutoff values had a very minimal effect of the resulting z-scores.

Evaluating all variants for the 4 assays took 5,929 GPU hours using 10 NVIDIA Tesla K20X GPUs.

### Integration with xQTLs

We downloaded QTLs for gene expression, DNase and histone modifications on Yoruban individuals (Grubert et al. 2015), QTLs for gene expression on LCLs from multiple European populations (Lappalainen et al. 2013). Enrichment was evaluated by comparing the DeepFIGV absolute z-cores from the lead QTLs to the scores for to variants ranked between 5^th^ and 10^th^. Statistical fine mapping results were obtained from multiple cell types (Brown et al. 2017). Rare variants associated with gene expression outliers from multiple tissues were obtained from GTEx (Li et al. 2017a). Enrichments are evaluated based on 2,113 rare variants associated with outliers and 67,044 not associated with outliers.

### Cancer somatic variants driving gene expression

We downloaded somatic variants in tumors that were identified by whole genome sequencing and results from an eQTL analysis combining nearby variants and testing the association with proximal genes (Zhang et al. 2018). We considered the 569 genes with cis-eQTLs at FDR < 30% and evaluated the DeepFIGV z-score for each of 4 epigenetic assays for the 2309 somatic variants in the proximal regions. The enrichment analysis compared these variants to somatic variants in this dataset that were not associated with gene expression changes and which were matched for distance to transcription start site.

### Prediction of allele specific binding (ASB)

DeepFIGV scores were used to predict the presence and direction of ASB using sites identified from transcription factor ChIP-seq and DNase I hypersensitivity experiments in LCLs (Chen et al. 2016b; Shi et al. 2016) and HeLa-S3 cells (Shi et al. 2016). For AlleleDB (Chen et al. 2016b), the ASB status for 42 ChIP-seq targets across 14 individuals totaling 77 experiments were reported at a total of 276,589 sites with sufficient read coverage (`accB.auto.v2.1.aug16.txt.gz`). Shi, et al. (Shi et al. 2016) reported ABS for 36 targets across 7 LCL and HeLa-S3 cells across 51,518 total sites (`ASB_GM12878_HeLa_1based.txt, ASB_other_GMs_1based.txt`). Sites from the two databases were combined to produce a non-redundant set, so sites identified for the same target and individual in both databases were not double counted. Sites were considered as ASB or non-ASB based on a Benjamini-Hochberg corrected p-value (beta-binomial for AlleleDB and binomial for Shi, et al.) < 0.05, or > 0.99, respectively. Only assays with at least 20 ASB examples were considered.

Precision-recall (PR) curves and area under the PR curve (AUPR) were used to evaluate the classification performance since the ASB vs non-ASB class counts were very imbalanced. PR curves and AUPR of empirical and random classes classifiers were evaluated with the PRROC package (Grau et al. 2015).

No allele-specific signal was used in the training of DeepFIGV and no re-training was performed for the ASB analysis. The DeepFIGV z-scores for each of the 4 assays in the training set were extracted for each site, and the PR and AUPR were computed by the intersecting these scores with the combined ASB dataset. Classifying ASB sites from non-ASB sites used the absolute value of the z-scores, while classifying the direction of ASB used the z-score itself. ASB magnitude was encoded as the number of alternative reads at a site divided by the total number of reads at that site. Thus, a positive ASB magnitude corresponds to a positive DeepFIGV z-score indicating that the alternative allele is predicted to increase signal compared to the reference allele.

### Computing disease enrichments: LD-score regression

Publically available GWAS summary statistics were obtained for immune diseases as well as other representative diseases and traits. We performed a partitioned hereditably analysis with LD-score regression (LDSC) (Finucane et al. 2015) in order to quantify the contribution to the trait heritability of variants with high absolute DeepFIGV z-scores. The per-SNP heritability was computed after accounting for the 32 other genomic annotations. Annotations included 28 provided with LDSC baseline model (i.e. TFBS, TSS, UTR, intron, promoter, enhancer, superenhancer, epigenetic assays multiple sources (H3K27ac, H3K4me1, H3K4me3, H3K9ac, DNase)) in addition to peak regions from the 4 assays in LCLs used by DeepFIGV. DeepFIGV was the only annotation with nucleotide level resolution; other annotations were 10s or 100s of bases wide.

This analysis was restricted to common variants outside of the MHC region. The per-SNP heritability was evaluated for site exceeding absolute z-score cutoffs for 1, 2, 3, 4 and 5. There we not a sufficient number of sites with larger scores, since only common variants were considered.

### Computing disease enrichments: Candidate causal variants

Candidate causal SNPs identified from finemapping analysis of autoimmune diseases were obtained from http://www.broadinstitute.org/pubs/finemapping (Farh et al. 2015). The DeepFIGV z-scores were obtained for the 8741 candidate causal SNPs for 39 traits. Enrichments for each trait were evaluated by comparing the number of candidate causal sites with a DeepFIGV absolute z-score exceeding a given cutoff to the expected value from a random set of sites from a null distribution. This null was constructed for each site by drawing 10,000 sites from across the genome matching the MAF, gene density, distance to nearest genes and number of sites within LD of 0.5 of the original site (Pers et al. 2015).

### Comparison to other variant scoring methods

Variant-level scores were obtained from DeepSea (Zhou and Troyanskaya 2015) evaluated on 17 DNase datasets and deltaSVM (Lee et al. 2015) evaluated on 35 DNase datasets, and CAPE (Alvarez et al. 2018) evaluated on 2 datasets. In addition we included CADD (Kircher et al. 2014) and LINSIGHT (Huang et al. 2017b). Scores were obtained from: https://www.ncbi.nlm.nih.gov/research/snpdelscore/rawdata/ For DeepSea and deltaSVM the reported delta values for the predicted signal from the reference and alternative alleles were transformed to a z-score using the observed standard deviation. Analysis was performed on a shared set of 12 million variants.

### Analysis of LCL MPRA results

Results were downloaded from Tewhey, et al. (2016) and variants were dived into 3 classes: 1) expression modulating variants that showed significant difference in expression between reference and alternative alleles, 2) variants that drove expression but did not show allelic differences, and 3) variants whose sequence did not drive expression in this assay. Enrichment of expression modulating variants (i.e. class 1) were compared to the other two classes as a function of high predicted epigenetic signal or DeepFIGV z-scores for each assay.

## Acknowledgements

We thank Yungil Kim for providing data on gene expression outliers. We thank Judy Cho and Kristen Brennand for feedback on the manuscript. This work was supported by NIMH 5R01MH109897. Gabriel E. Hoffman is partially supported by a NARSAD Young Investigator Award from the Brain and Behavior Research Foundation. This work was supported in part through the computational resources and staff expertise provided by Scientific Computing at the Icahn School of Medicine at Mount Sinai. We particularly thank Gene Fluder and Hyung Min Cho for assistance with GPUs.

## SUPPLEMENTARY FIGURES

**Supplementary Figure 1:**
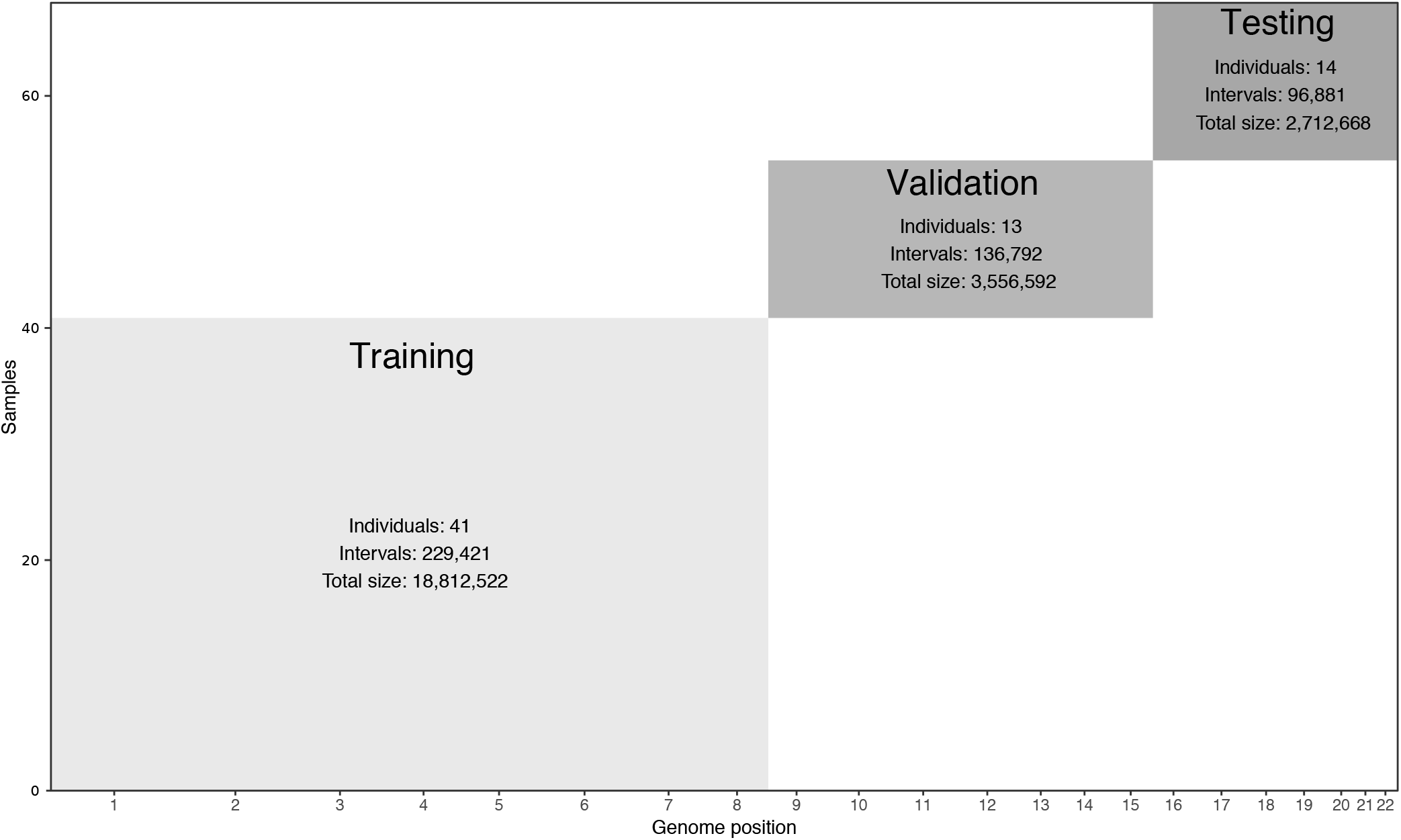
Design of training, validation and testing sets for DNase. The training set is composed of peaks from 41 individuals on chromosomes 1-9. The validation set is composed of peaks from 13 individuals on chromosomes 8-16. The test set is composed of peaks from 14 individuals on chromosomes 16-22.

**Supplementary Figure 2:**
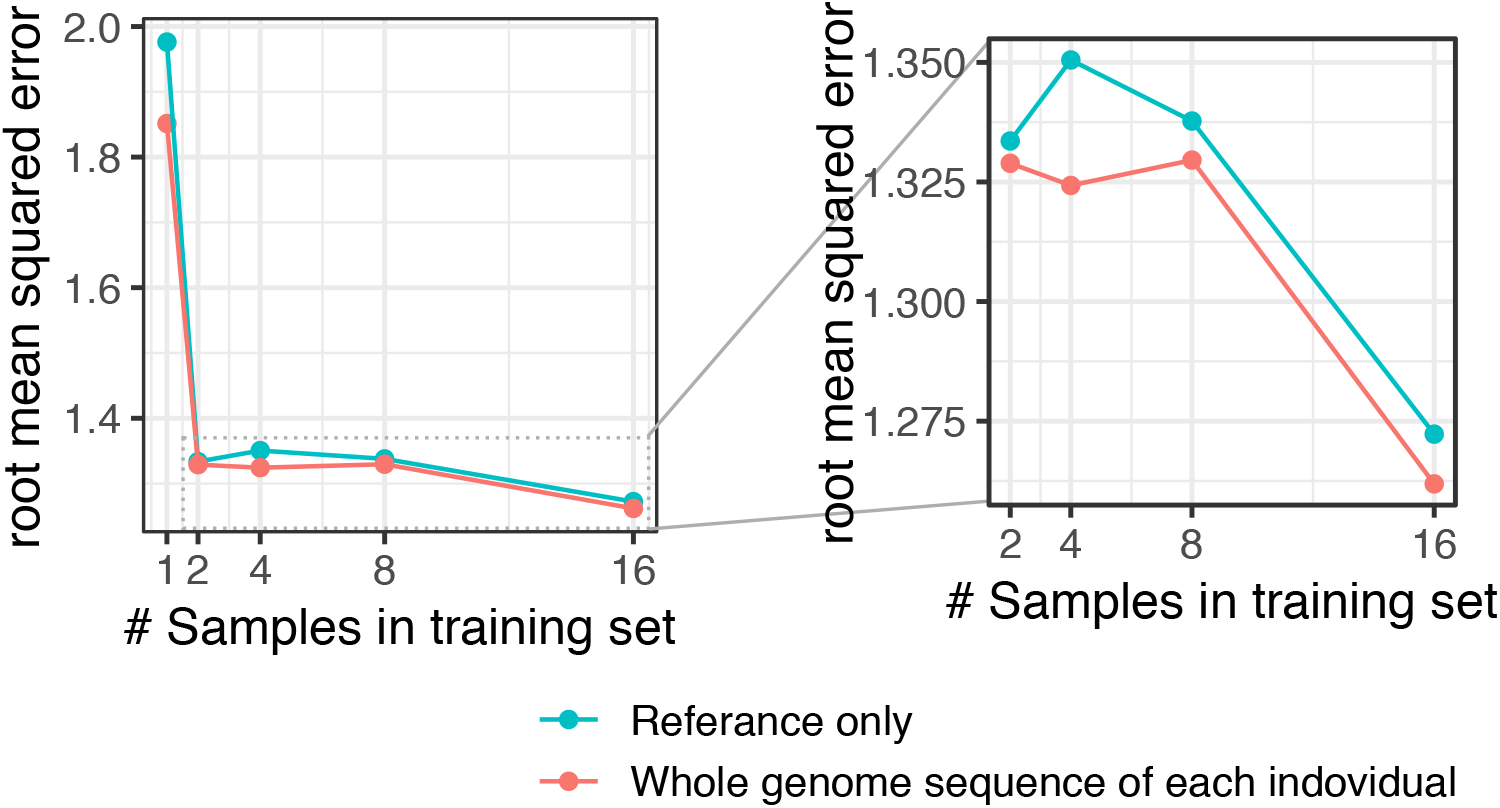
Design of training dataset decreases prediction error. Increasing the number of individuals in the training set and including genetic variation from whole genome sequencing decreased the prediction error for DNase signal. Left panel shows results for training on between 1 and 16 individuals. Right panel is the same results but zoomed in.

**Supplementary Figure 3:**
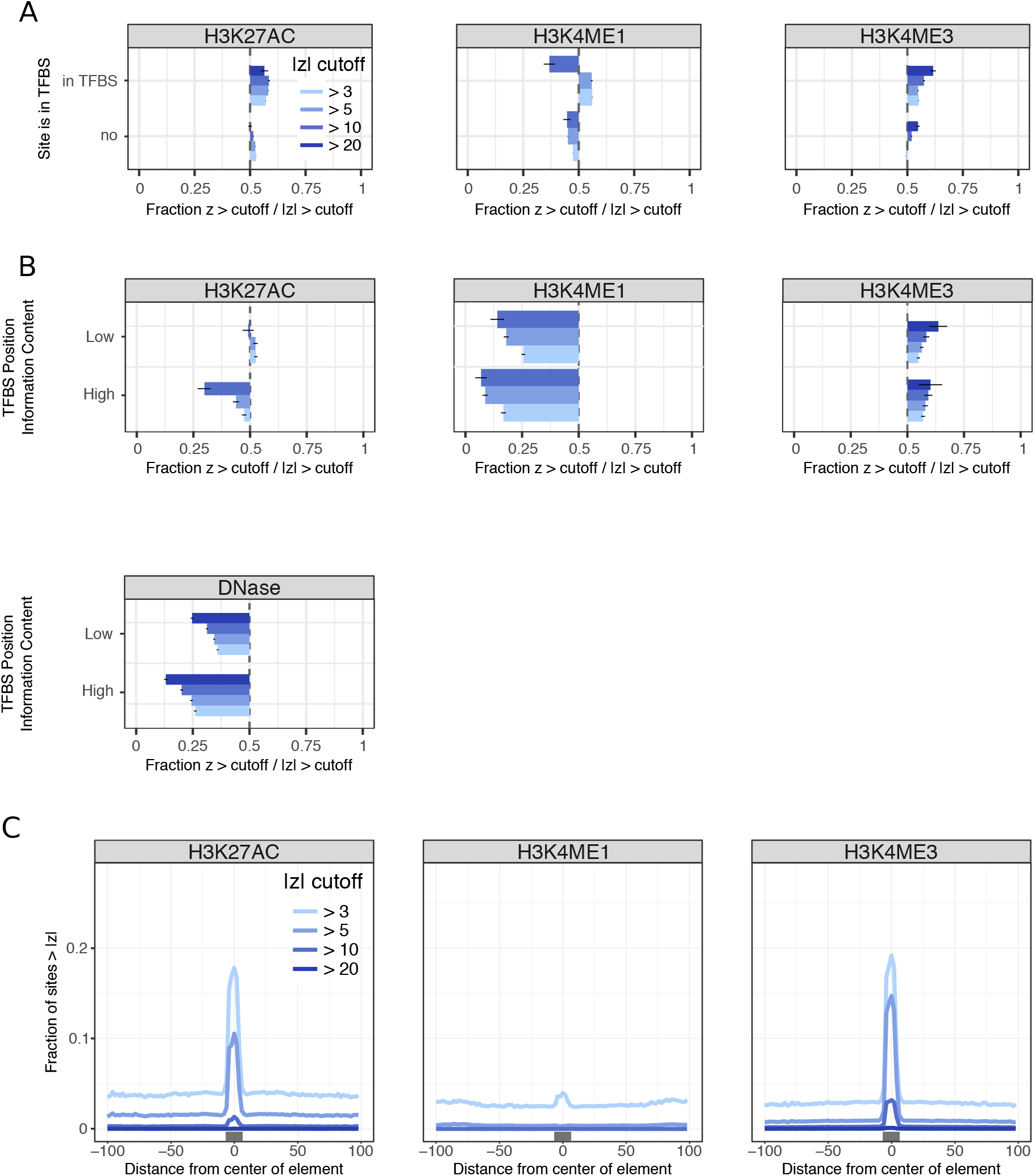
Transcription factor binding site motifs are enriched for variants with large DeepFIGV absolute z-scores. **A)** Ratio indicating the fraction of sites with positive DeepFIGV z-scores for 4 cutoffs. Ratios are shown for sites with a TFBS motif or not in a TFBS motif. A value of 0.5 indicates an equal number of variants with positive and negative z-scores. A value < 0.5 indicates an excess of variants with negative scores that decrease the epigenetic signal. **B)** Ratio indicating the fraction of sites with positive DeepFIGV z-scores for sites in TFBS motifs annotated where each site is annotated as low or high information content. **C)** Fraction of sites exceeding 4 cutoffs for DeepFIGV absolute z-score based on distance to TFBS motifs. Black box indicates median size of TFBS motif.

**Supplementary Figure 4:**
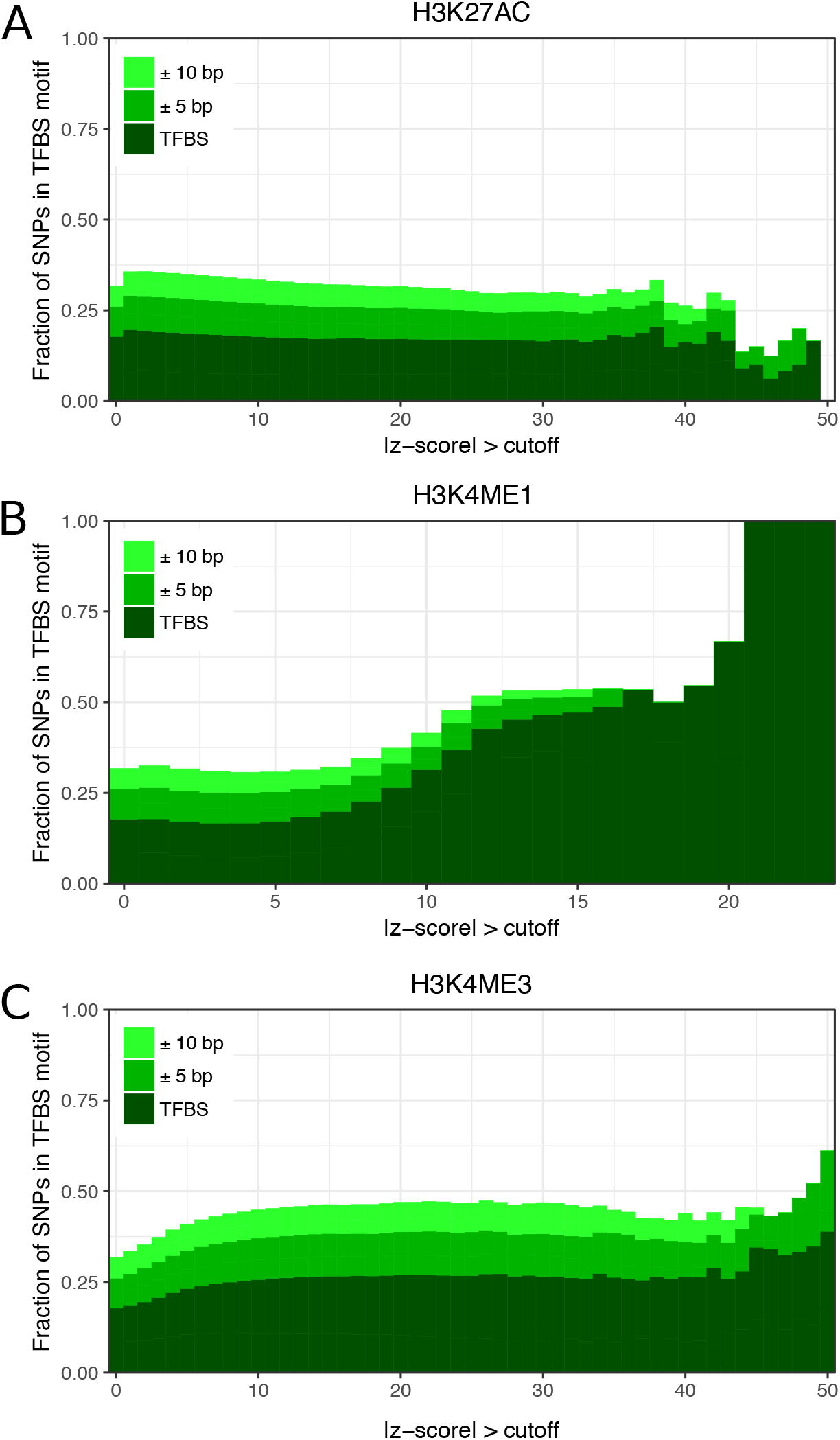
Fraction of sites that are in a transcription factor binding site motif, or in the flanking 5 or 10 bp, for a range of DeepFIGV absolute z-score cutoffs. Results are shown for for **A)** H3K27AC, **B)** H3K4ME1 and **C)** H3K4ME3.

**Supplementary Figure 5:**
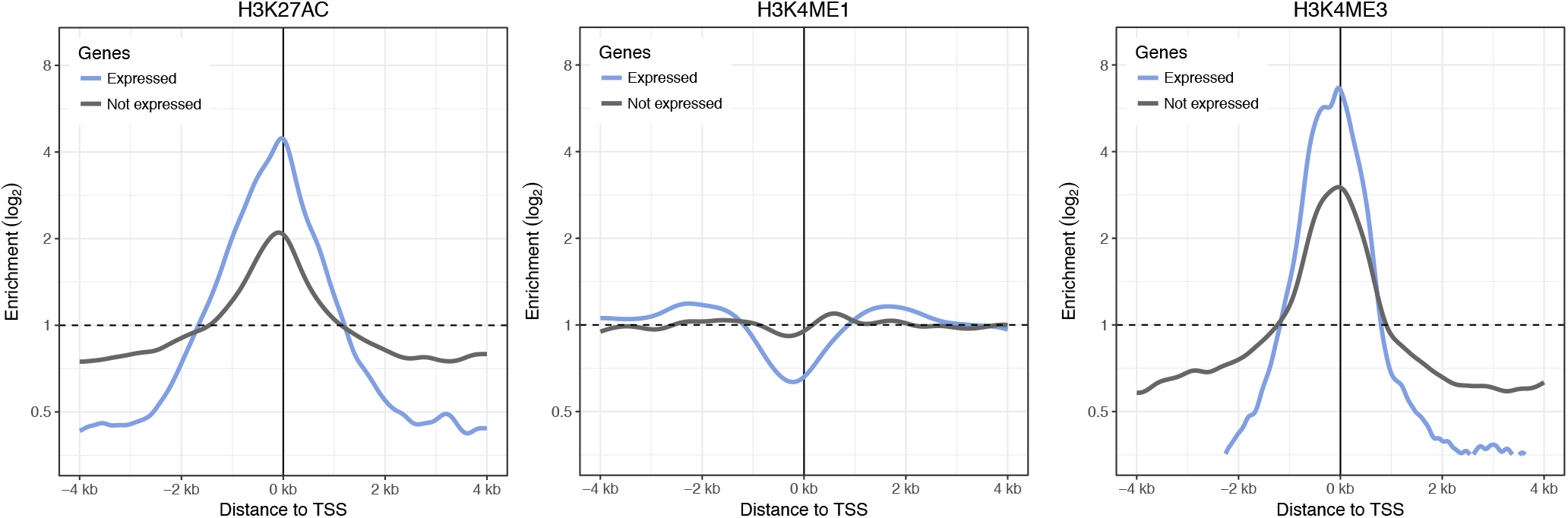
Enrichment of variants with large DeepFIGV absolute z-scores around transcription start sites. Enrichment of sites with absolute z-scores greater than 5 near the transcription start site of genes stratified by whether the genes are expressed in LCLs. Sites with absolute z-scores less than the genome-wide mean are used as the baseline for the enrichment.

**Supplementary Figure 6:**
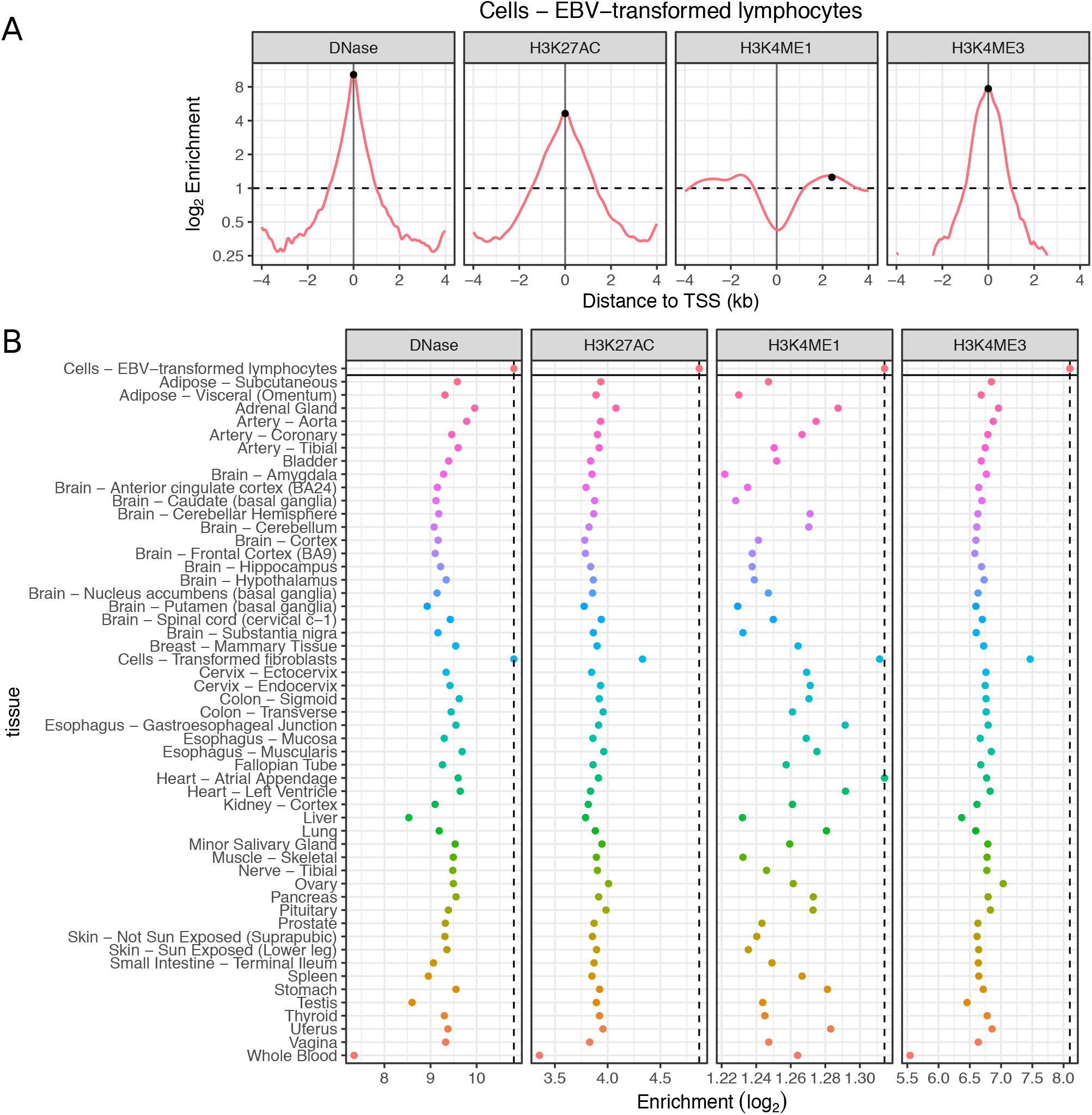
Enrichment around transcription start sites of tissue-specific genes. For each of 53 tissues from GTEx, the expression magnitude of each gene was summarized as the mean transcripts per million (TPM) from all samples from that tissue. Tissue-specific genes were defined by evaluating the set of protein coding genes expressed in each GTEx tissue and subtracting the set of ubiquitously expressed genes. An expression cutoff of >= 1 TPM was used. Ubiquitously expressed genes were determined based on passing this cutoff in all 53 tissues. **A)** The enrichment around the transcription start site of the LCL-specific genes was evaluated for variants with |z| > 5 compared to variants with |z| less than the genome-wide mean. Enrichments are show for variants from 4 epigenetic assays. Black circles indicate the maximum enrichment value used. **B)** The maximum enrichment value is shown for each tissue and assay. For LCL-specific genes, this value corresponds to the largest enrichment for each assay as shown in **(A).** The values for all other genes are defined based on the set of tissue-specific genes. The vertical dashed line indicates the enrichment for LCL-specific genes. Since the DeepFIGV model was trained on LCLs (but from a different dataset), the fact that LCL-specific genes show the largest enrichment illustrates the specificity of the DeepFIGV variant impact scores.

**Supplementary Figure 7:**
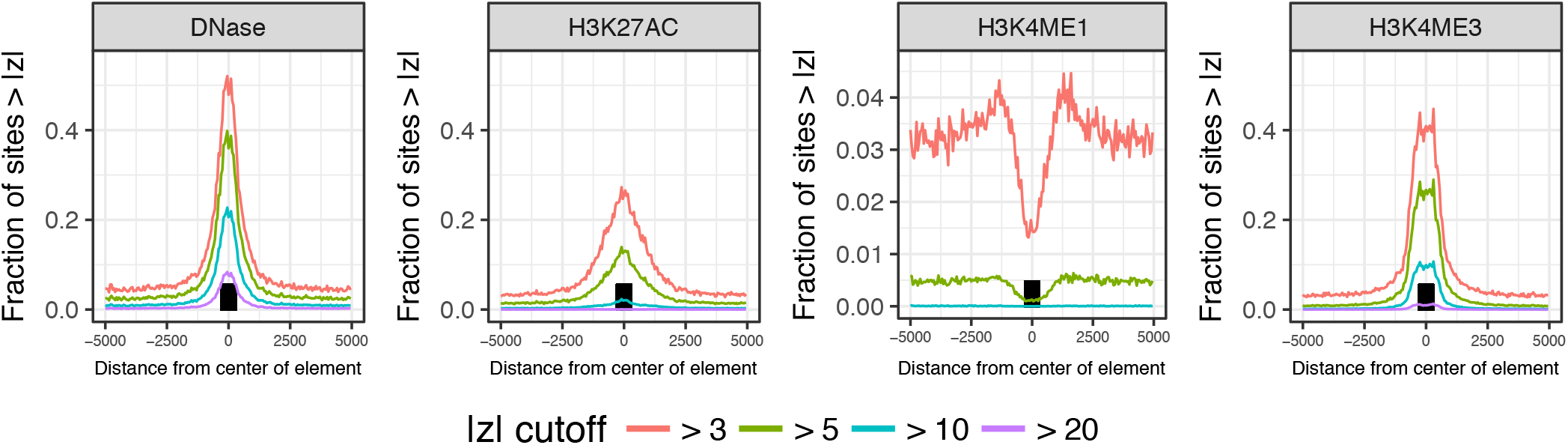
Enrichment of variants with large DeepFIGV absolute z-scores in CpG islands. Fraction of sites exceeding 4 cutoffs for DeepFIGV absolute z-score based on distance to CpG islands. Results are shown for DeepFIGV z-score for 4 epigenetic assays. Black box indicates median size of CpG island. That the H3K4ME1 is shown in a different scale.

**Supplementary Figure 8:**
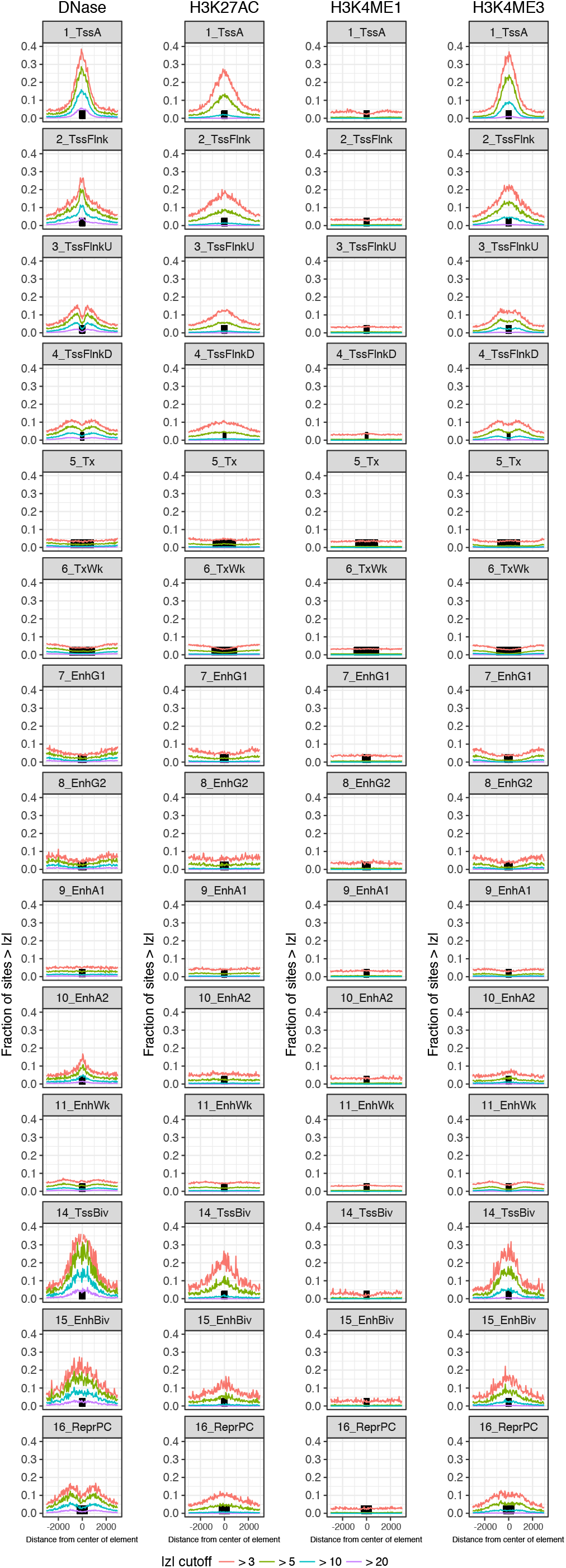
Enrichment of variants with large DeepFIGV absolute z-scores in ChromHMM segments. Fraction of sites exceeding 4 cutoffs for DeepFIGV absolute z-score based on distance to ChromHMM segment from LCLs (GM12878). Results are shown for DeepFIGV z-score for 4 epigenetic assays. Black box indicates median size of ChromHMM segment, if it is less than 5kb. ChromHMM tracks with no enrichment are omitted.

**Supplementary Figure 9:**
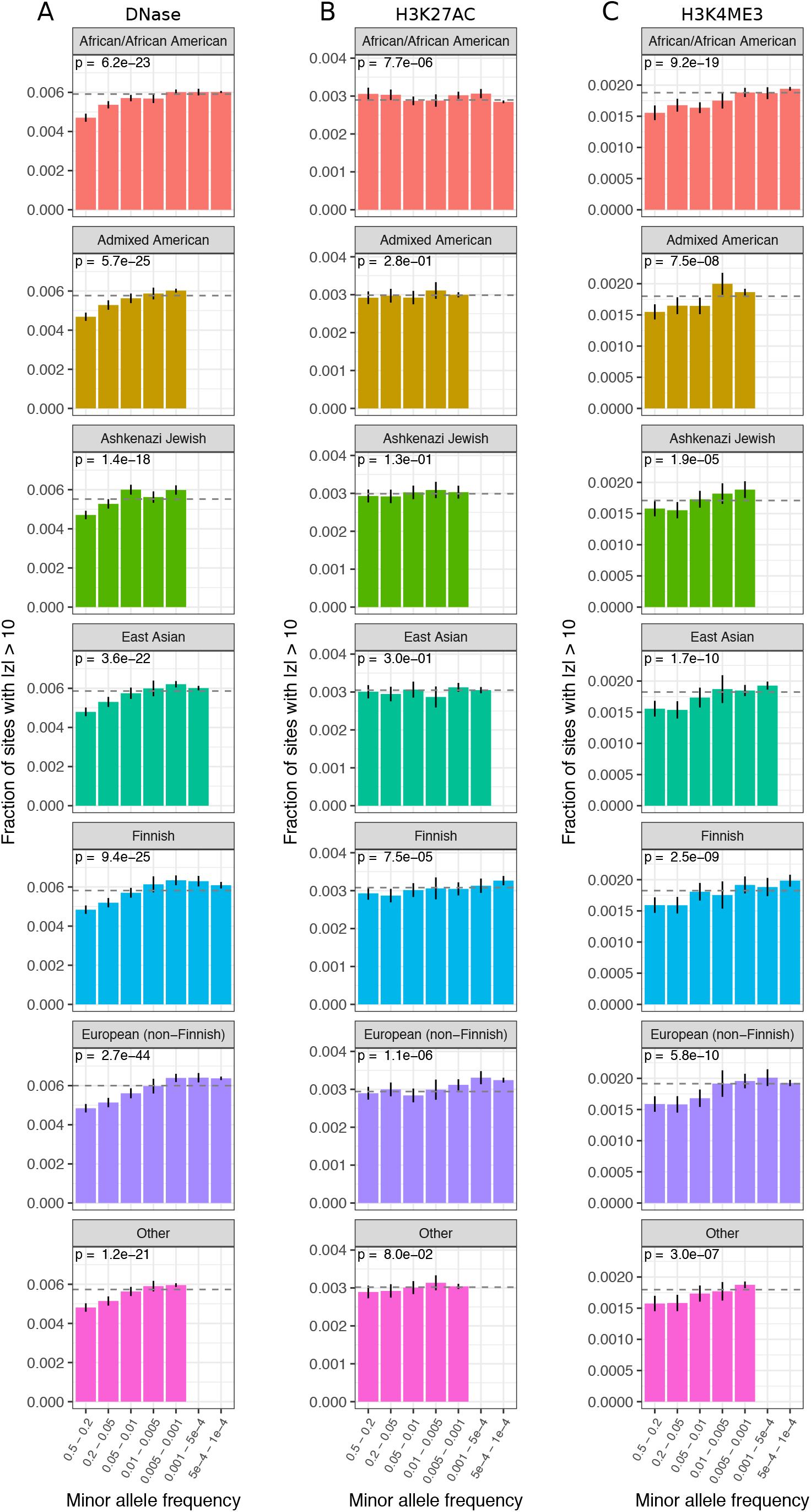
Relationship of predicted functional score to allele frequency. Fraction of sites with absolute z-scores for **A)** DNase, **B)** H3K27ac, and **C)** H3K4me3 greater than 10 within 7 minor allele frequency bins based on 7 populations from gnomAD. Dashed line indicates genome wide fraction of sites. P-value is based a logistic regression where the response is a binary variable indicating if the absolute z-scores is greater than 10 and the log minor allele frequency is the predictor. Error bars show 95% confidence intervals.

**Supplementary Figure 10:**
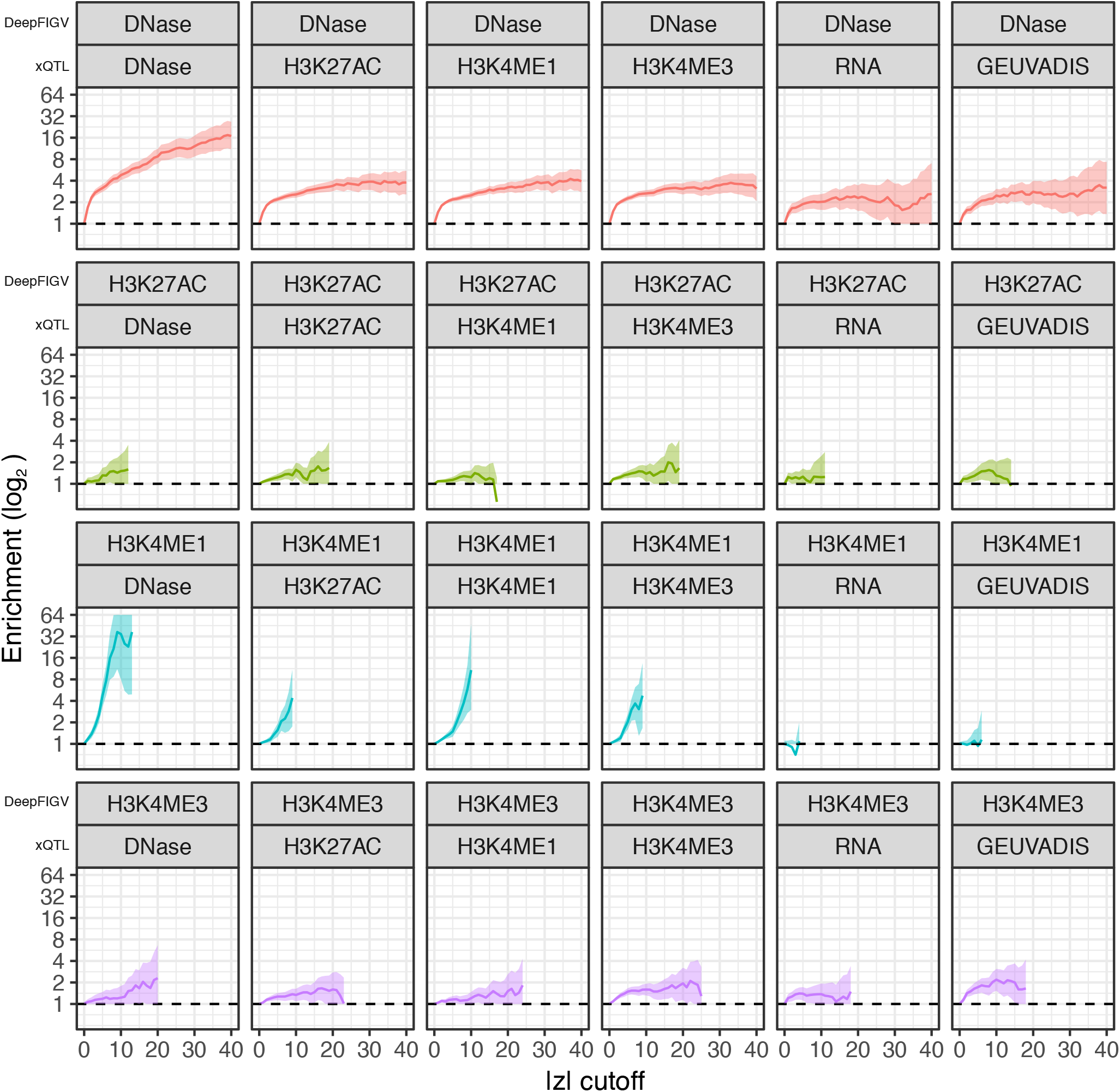
Lead xQTL variants are enriched for large DeepFIGV absolute z-scores. Lead variants from xQTL analysis from lymphoblastoid cell lines are enriched for SNPs with DeepFIGV absolute z-score exceeding a range of cutoffs compared to variants ranked between 5^th^ and 10^th^. Enrichments are evaluated using DeepFIGV scores for 4 assays for QTLs DNase, H3K27ac, H3K4me1, H3K4me3, and gene expression (i.e. RNA). from 69 Yoruban individuals (Grubert et al. 2015). The last column indicates expression QTLs from 373 European individuals from the GEUAVIDIS study (Lappalainen et al. 2013). Shaded regions indicate 95% confidence intervals.

**Supplementary Figure 11:**
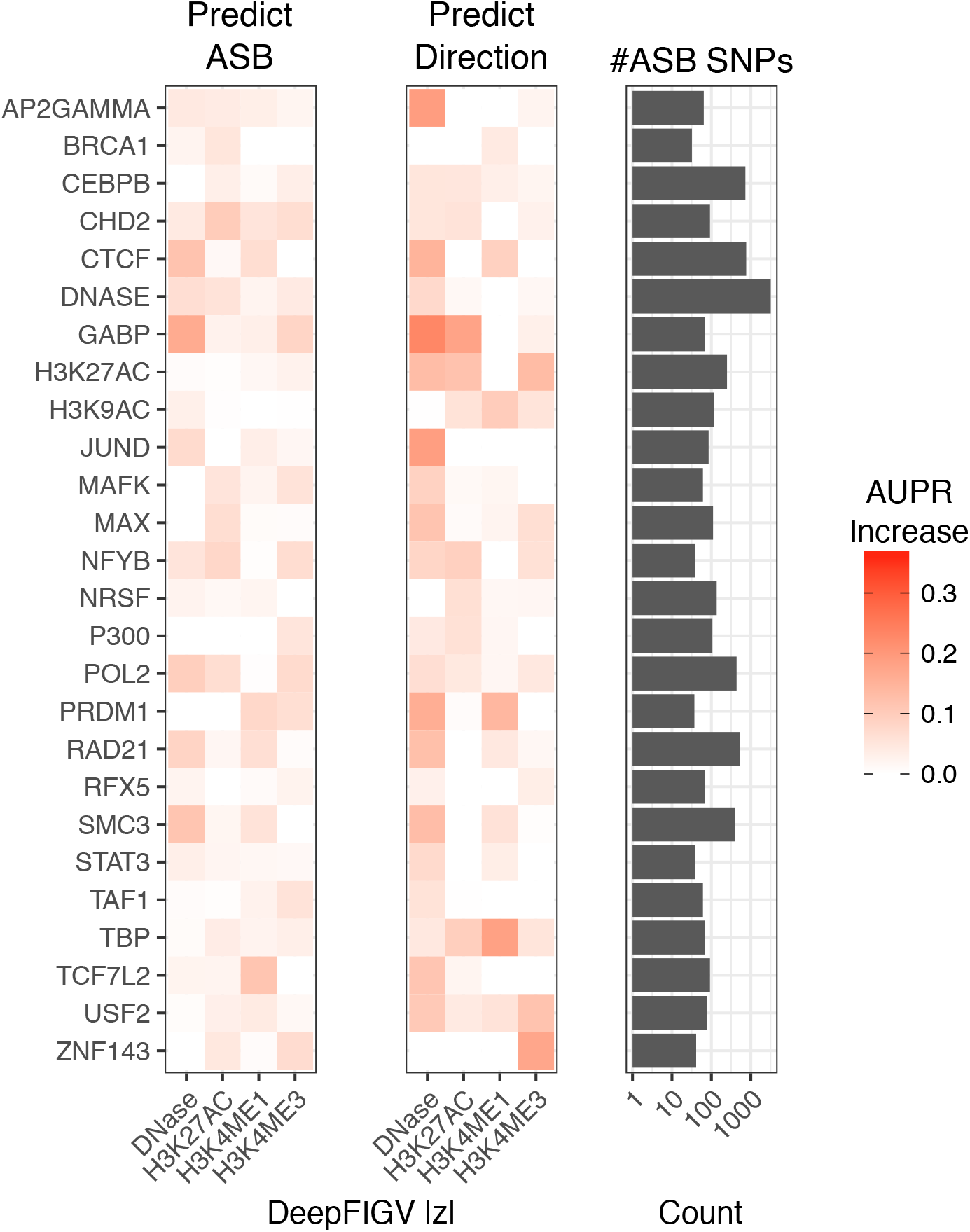
DeepFIGV scores predict allele specific transcription factor binding in HeLa-S3 cells. Increase in AUPR of predicting ASB status for DeepFIGV scores for 4 epigenetic assays compared to a random predictor. Increase in AUPR is shown for predicting ASB versus no ASB (left) and predicting the directionality of ASB (reference versus alternative) (center). Right panel shows the number of ASB SNPs considered in each analysis.

**Supplementary Figure 12:**
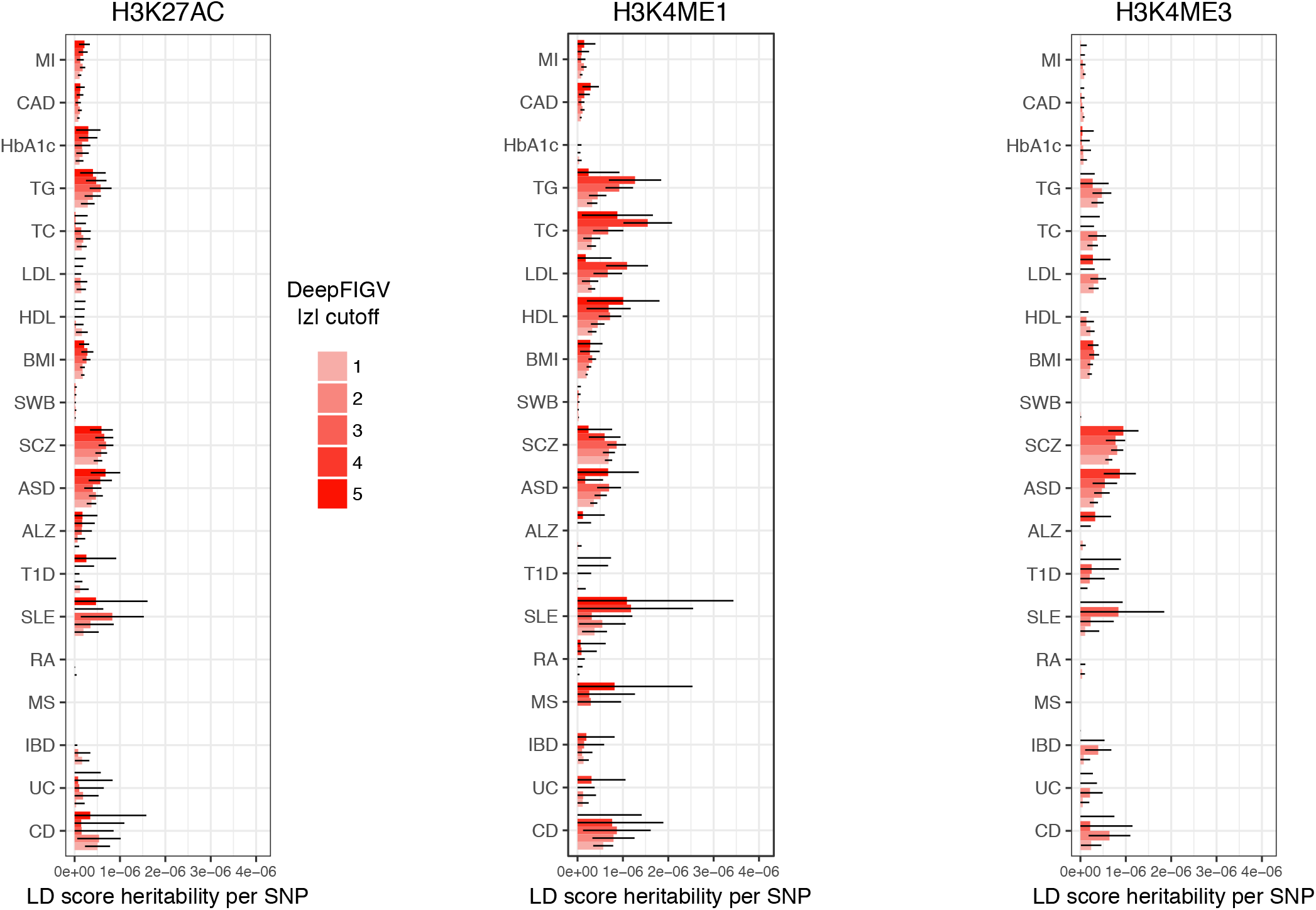
Disease risk variants are enriched for large DeepFIGV scores for partitioned heritability analysis. Linkage-disequilibrium score regression partitioned heritability estimates for 3 epigenetic assays: for **A)** H3K27ac, **B)** H3K4me1 and **C)** H3K4me3. Heritability per SNP is computed for variants exceed 5 cutoffs for DeepFIGV absolute z-score for each assay. Error bars indicate 2 standard deviations.

**Supplementary Figure 13:**
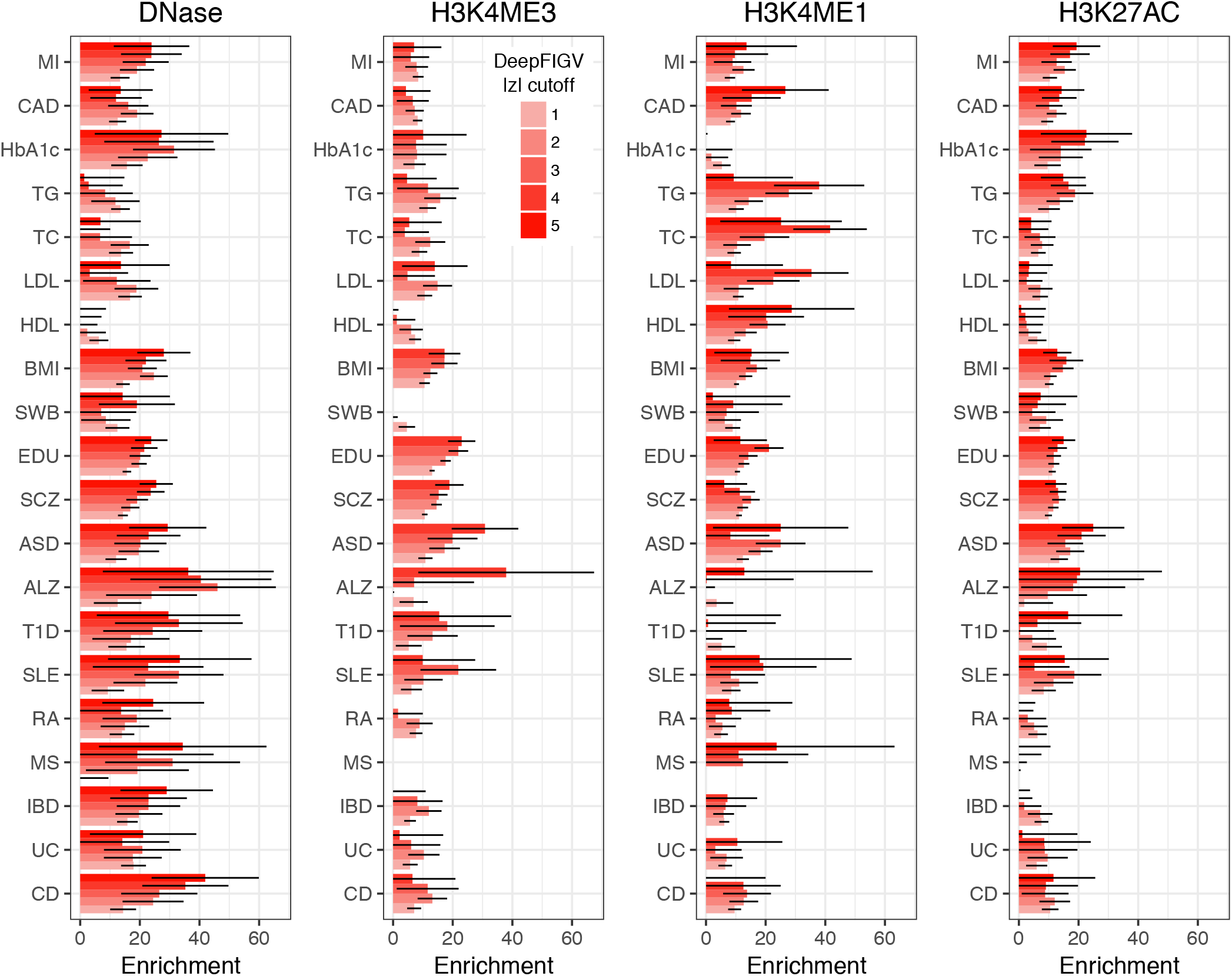
Heritability enrichment from linkage-disequilibrium score regression. Estimated enrichment in heritability from same analysis as in Supplementary Figure 10.

**Supplementary Figure 14:**
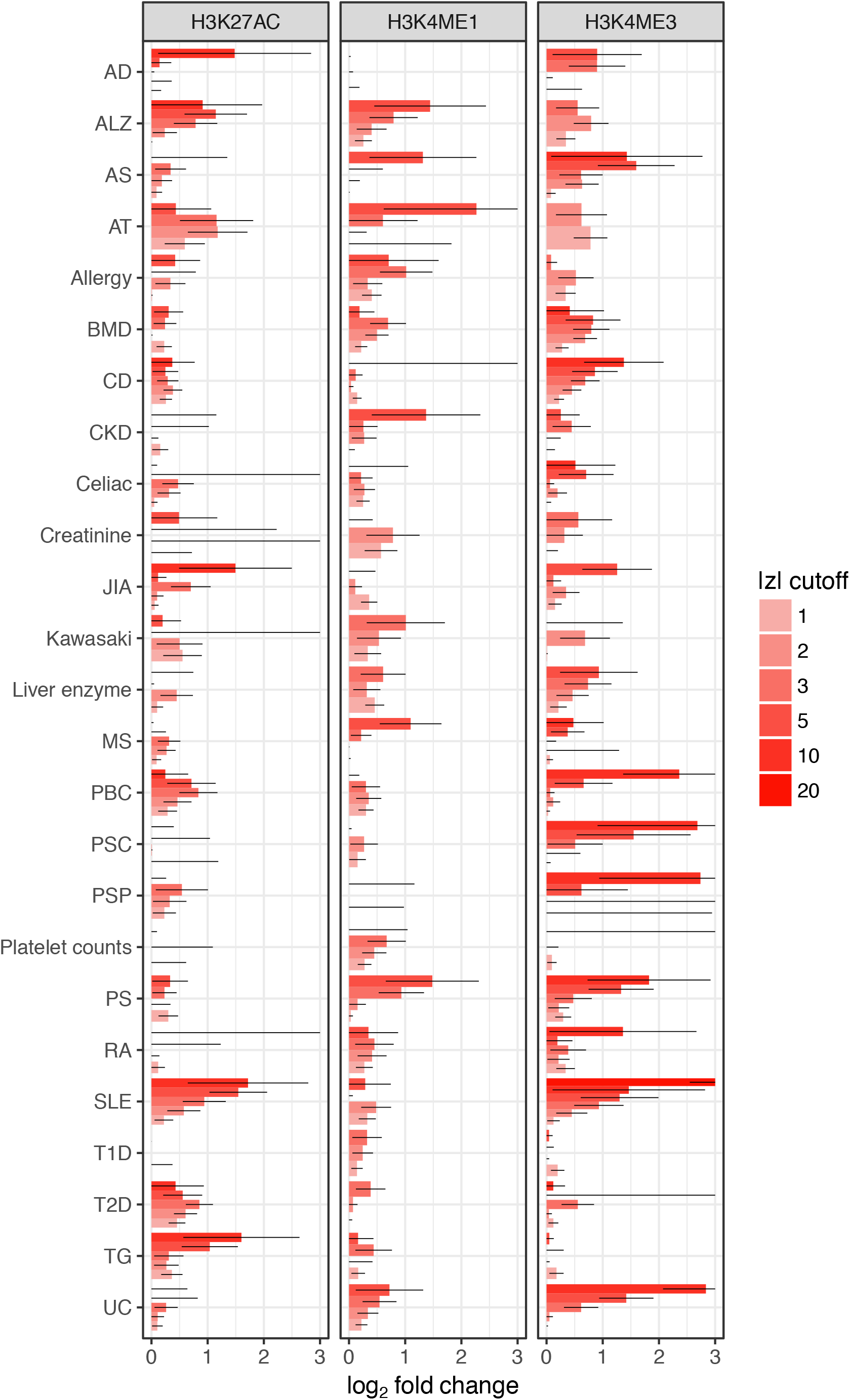
Enrichment of candidate causal variants for autoimmune disease. Enrichments are evaluated for variants exceeding 6 cutoffs DeepFIGV absolute z-score for H3K27ac, H3K4me1 and H3K4me3. Error bars indicate 2 standard deviations. Disease abbreviations: AD (Atopic dermatitis), ALZ (Alzheimer’s), AS (Ankylosing spondylitis), ASD (Autism spectrum disorder), AT (Autoimmune thyroiditis), BMD (Bone mineral density), BMI (Body mass index), CAD (Coronary artery disease), CD (Crohn’s disease), CKD (Chronic kidney disease) HbA1c (HbA1c protein level in blood), HDL (High-density lipoprotein), IBD (Inflammatory bowel disease), JIA (Juvenile idiopathic arthritis), LDL (Low-density lipoprotein), Liver enz (gamma glutamyl transferase), MI (myocardial infarction), MS (Multiple sclerosis), PBC (Primary biliary cirrhosis), PSC (Primary sclerosing cholangitis), PSP (Progressive supranuclear palsy), PS (Psoriasis), RA (Rheumatoid arthritis), SLE (Systemic lupus erythematosus), SWB (Subjective well-being), T1D (Type 1 diabetes), T2D (Type 2 diabetes), TC (total cholesterol), TG (Triglycerides), UC (Ulcerative colitis).

**Supplementary Figure 15.**
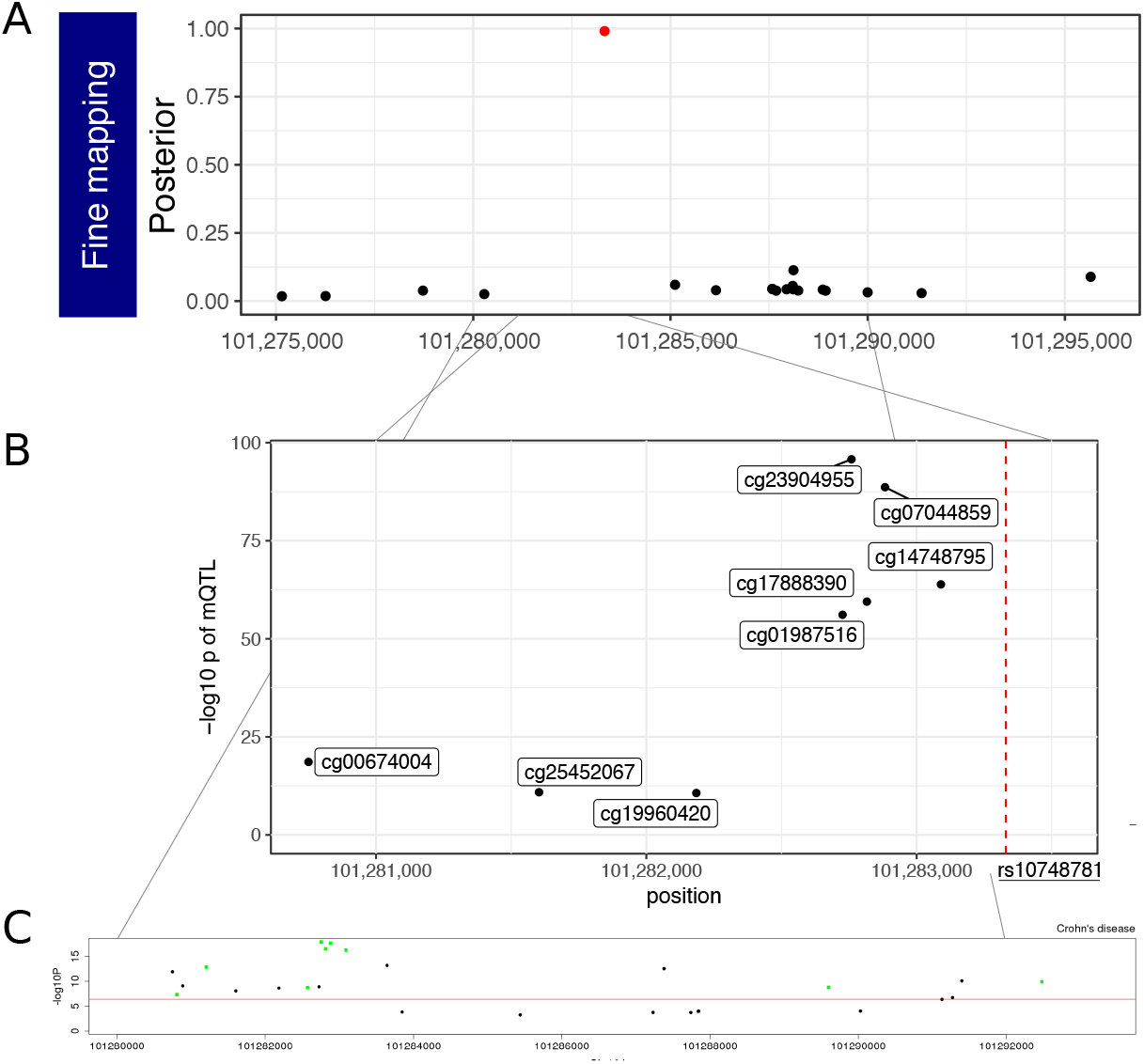
Inflammatory bowel disease risk from rs10748781 is mediated by DNA methylation. **A)** Posterior probability of each variant in chr10:101,275,149-101,295,862 being causal. Red point indicates rs10748781. **B)** rs10748781, shown as a red line, is a methylation QTL for CpG sites in this region (Hannon et al. 2017). **C)** Summary Mendelian randomization analysis of inflammatory bowels disease where each point is a DNA methylation probe.

**Supplementary Figure 16.**
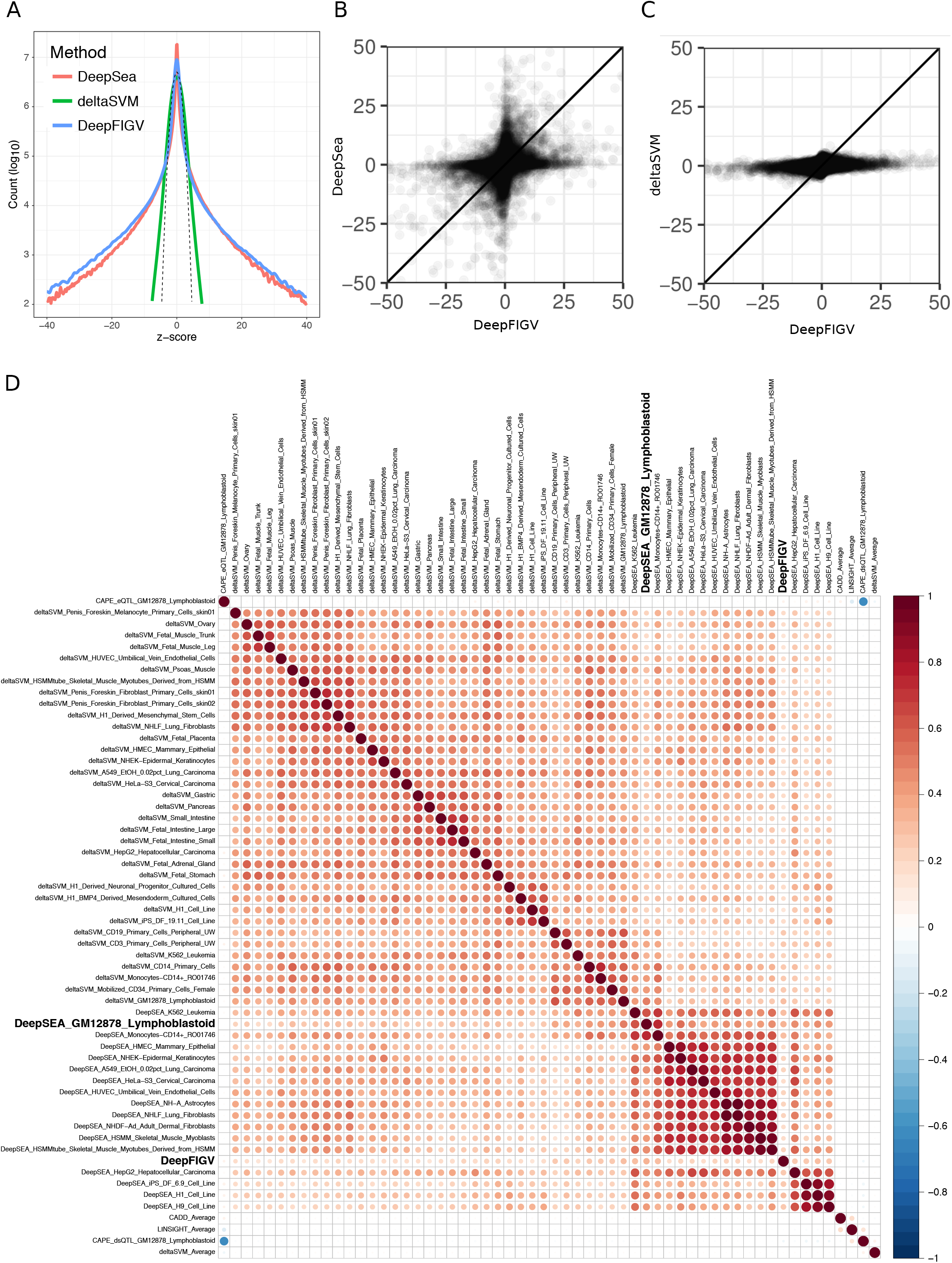
Comparison of DeepFIGV scores for DNase with DNase scores from other methods. All methods were evaluated on a shared set of 12 million variants. **A)** Density plot of z-scores for DeepFIGV as well as DeepSea (Zhou and Troyanskaya 2015) and deltaSVM (Lee et al. 2015) evaluated on DNase data from LCL GM12878. Dashed line indicates the null distribution of the z-scores, which is the standard normal distribution. **B)** Plot of z-scores from DeepFIGV and DeepSea. **C)** Plot of z-scores for DeepFIGV and deltaSVM. **D)** Spearman correlation between all pairs of XXX scores. Scores include DeepFIGV, plus DeepSea and deltaSVM evaluated on data from multiple cell types. Also included are CAPE (Li et al. 2017b), CADD (Kircher et al. 2014) and LINSIGHT (Huang et al. 2017b) methods.

**Supplementary Figure 17.**
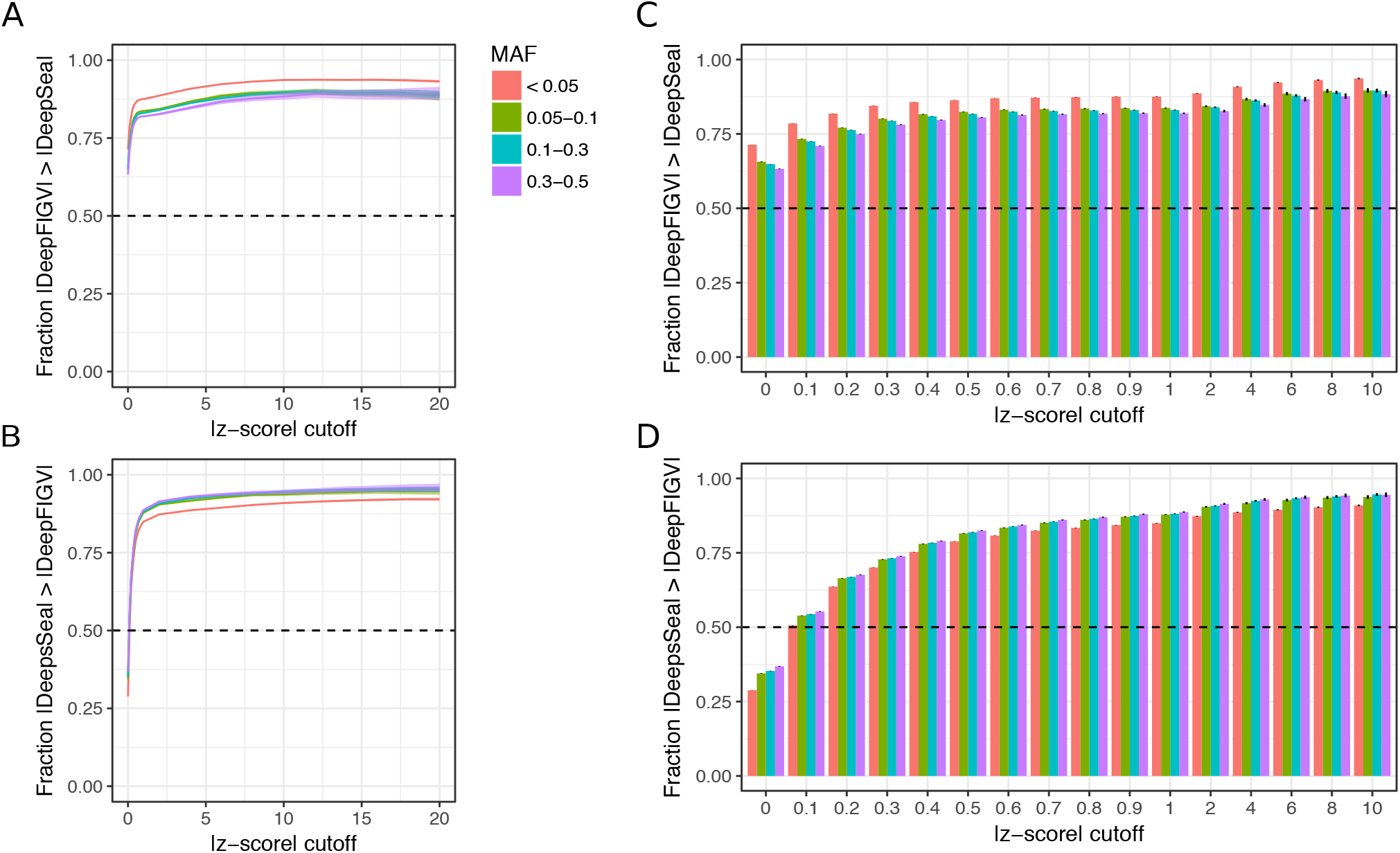
Comparison between DeepFIGV and DeepSea (evaluated on LCL GM12878) as a function of allele frequency. **A,B)** Evaluate the fraction of variants for which the z-score from one methods exceeds the z-score from the other method. For variants that pass a z-score cutoff from the first method, plot the fraction of sites for which the z-score from the first method is greater than the z-score from the second method. Fractions are shown for a range of z-score cutoffs and are stratified by the minor allele frequency in the Yoruban population. Fractions were evaluated by a applying the z-score cutoff to **A)** DeepFIGV and **B)** DeepSea. **C,D)** Barplots show finer detail from the plots in **(A)** and **(B).** Fractions were evaluated by a applying the z-score cutoff to **C)** DeepFIGV and **D)** DeepSea. Error bars indicated 2 standard deviations.

**Supplementary Figure 18.**
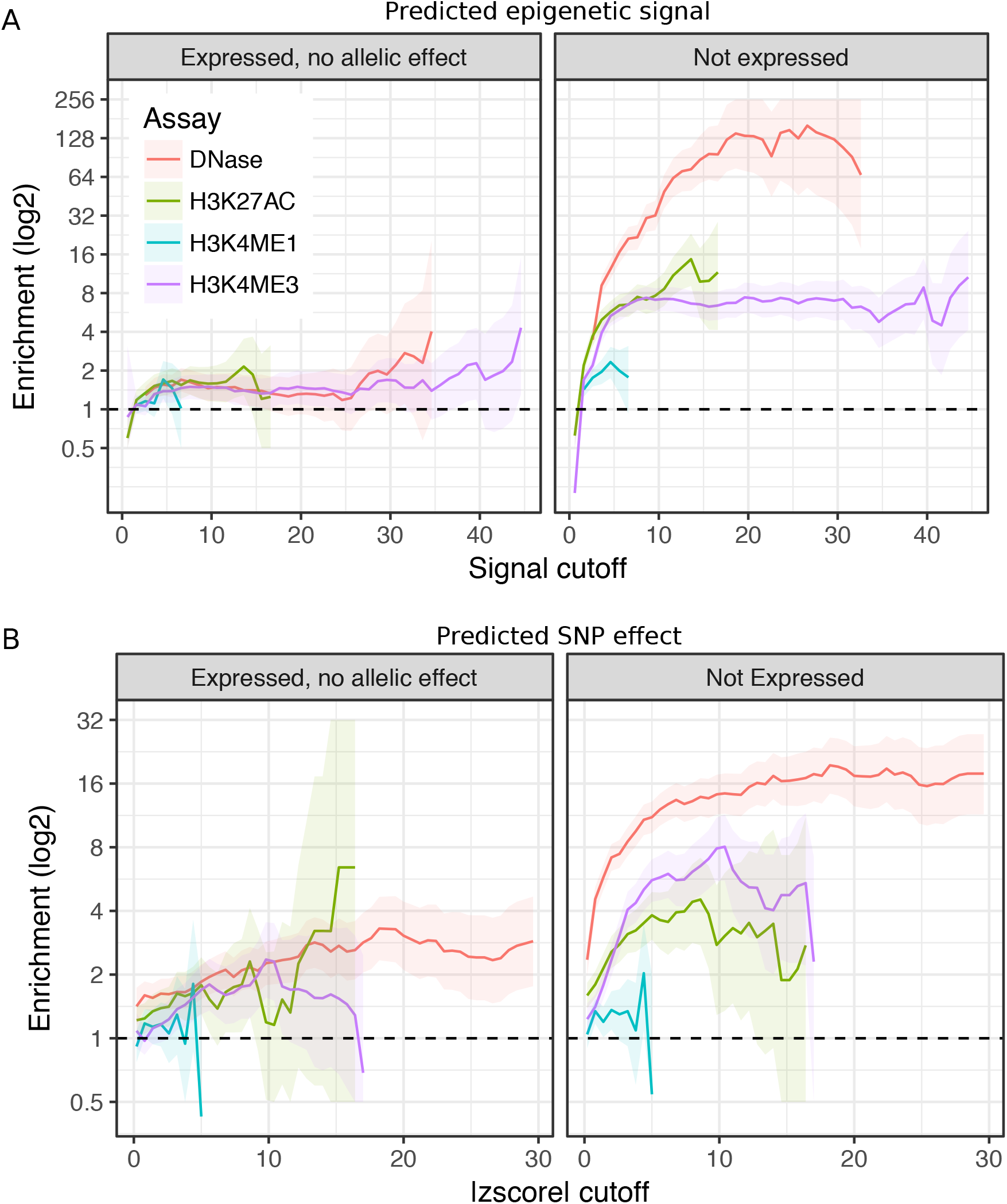
Enrichment of analysis of DeepFIGV with experimental massively parallel reporting assay (MPRA). Tewhey, et al. performed an MPRA of 32K variants in LCLs by inserting 150 bp sequences centered at the variant into an episomal vector (Tewhey et al. 2016). Based on experimental readout, Tewhey, et al. divided the sequences into 3 classes: 1) expression modulating variants that showed significant difference in expression between reference and alternative alleles, 2) variants that drove expression but did not show allelic differences, and 3) variants whose sequence did not drive expression in this assay. Here, we compare the properties of variants with allelic effect (class 1) to variants that are expressed (class 2) or not expressed (class 3). **A**) DeepFIGV computational predictions of epigenetic signal distinguish sequences with expression modulating variants from sequences that do not drive expression. Expression modulating variants are enriched for having a strong epigenetic signal by multiple assays compared to variants that did not drive expression. Enrichment is shown for increasing epigenetic signal values. **B**) DeepFIGV predicted SNP effects can distinguish sequences with expression modulating variants from sequences with no allelic effect. Enrichment is shown for increase absolute z-score cutoffs. The enrichment compared to sequences that do not drive expression is largest. Distinguishing between sequence with and without allelic effects is more challenging because the experimental effect sizes are small (Tewhey et al. 2016), yet DNase predicted variant effects shown a significant enrichment across a range of cutoffs. Shaded regions indicate 95% confidence intervals.

**Supplementary Table 1. Summary of DNA sequence inputs to neural network and split of training, validation and test sets.**

**Supplementary Table 2. List of databases with variant sets**

**Supplementary Table 3. Parameters for basset convolutional neural network**

